# Gene loss, repression, amplification, and horizontal acquisition shape galactose/melibiose metabolism in fission yeast

**DOI:** 10.64898/2026.03.02.708787

**Authors:** Xiao-Min Du, Fang Suo, Li-Lin Du

## Abstract

Natural variation in metabolism is a key driver of microbial adaptation. While galactose utilization is well-studied in budding yeasts, it remains poorly understood in the fission yeast *Schizosaccharomyces pombe*. Here, we reveal extensive natural variation in galactose utilization across *S. pombe* isolates—from complete deficiency (Gal^−^) to exceptionally fast growth (Gal^F^). Gal^−^ strains fall into two classes: one with deletions of the *gal* gene cluster (via three distinct mechanisms), and another with intact but repressed *gal* genes. In contrast, Gal^F^ is driven by an amplified gene cluster absent from the reference genome—the *gal-mel* cluster (GMC)—which also confers melibiose utilization (Mel^+^). Mel^+^ is exclusively linked to the GMC, except in one strain harboring a standalone melibiase gene. Phylogenetic analyses indicate that horizontal gene transfer may underlie these adaptive traits. Together, our work demonstrates how diverse mechanisms—gene loss, repression, amplification, and horizontal acquisition—shape metabolic diversity and ecological specialization in fission yeast.

**Significance Statement:** Evolutionary adaptation is often viewed as a one-way street for a given trait, with a species either losing or gaining a function. Our results indicate this view is incomplete. We show that, for a single metabolic trait, a eukaryotic species can simultaneously pursue opposing evolutionary trajectories. Within *S. pombe*, reductive evolution through gene loss and repression occurs alongside expansive innovation via horizontal gene transfer and amplification. This bidirectional evolution reveals that adaptive potential is not confined to a single path but encompasses multiple concurrent strategies within a species’ gene pool. These findings support a refined model of eukaryotic genome plasticity, in which opposing evolutionary forces act in concert to generate a dynamic repertoire of metabolic capabilities.

## Introduction

Natural variation in microbes’ ability and efficiency in utilizing different nutrients underpins evolutionary adaptation to shifting environments (1–3). Understanding the mechanisms that generate this metabolic diversity is essential for comprehending microbial ecology, reconstructing evolutionary trajectories, and harnessing microbial potential in biotechnology.

The galactose utilization pathway in budding yeasts is a well-established model for studying the evolution of metabolic traits (4–12). In *Saccharomyces cerevisiae*, galactose utilization requires the transporter Gal2 and three Leloir pathway enzymes—Gal1, Gal7, and Gal10 (13–15). The genes encoding the Leloir enzymes are often clustered—a feature observed in diverse fungi including *Sac. cerevisiae*, *Candida albicans*, *Schizosaccharomyces pombe*, and *Cryptococcus neoformans* (4,5,9,16,17). Some *Torulaspora* strains even harbor expanded clusters encoding additional genes for galactose and melibiose utilization (7).

*S. pombe*, a well-known model organism in genetics and evolutionary biology (18–29), belongs to *Taphrinomycotina*—a lineage of *Ascomycota* distantly related to budding yeasts. It is globally distributed and frequently isolated from human-associated niches such as honey, dried fruits, grape mash, and various fermented beverages (29,30). Population genomic studies of 161 *S. pombe* natural isolates defined 57 non-clonal strains (24), later shown to descend from two ancient clades: REF and NONREF (21,25).

In *S. pombe* and other fission yeast species, the gene cluster containing the Leloir pathway genes—hereafter referred to as the *gal* cluster—was acquired via an ancient horizontal gene transfer (HGT) event from a budding yeast donor in the order *Serinales* (formerly known as the CUG-Ser1 clade) (5,9,31). This cluster contains *gal1*, *gal7*, and *gal10*, plus *SPBPB2B2.11*—a gene homologous to *C. albicans GAL102*, encoding a UDP-glucose 4,6-dehydratase unrelated to galactose metabolism (32). *SPBPB2B2.11* has been designated *tgd1* in PomBase (33). We propose renaming it *gal102* to reflect its HGT origin and use this designation hereafter.

Despite possessing galactose utilization genes, *S. pombe* is described in taxonomic literature as a species that cannot utilize galactose (34). Matsuzawa et al. showed that while wild-type *S. pombe* reference background strains cannot grow in liquid galactose media, a mutant strain that upregulates the expression of the *gal* cluster genes can grow (35). This indicates that the *S. pombe* reference background retains latent galactose utilization capacity but may require an induction signal to express this ability. It remains unclear to what extent the *S. pombe* reference background is representative of the species.

In this study, we identify ethanol as an induction signal that enhances the galactose utilization ability of the *S. pombe* reference background. Systematic analysis of *S. pombe* natural isolates reveals extensive phenotypic variation: some strains lack galactose utilization ability due to gene loss or repression, while others gain enhanced capability through a high-copy-number *gal-mel* cluster (GMC), which is absent in the reference genome. The GMC includes a melibiase gene enabling growth on melibiose (Mel^+^). Strain JB875 is also Mel^+^, due to a distinct, non-clustered melibiase gene; phylogenetic analyses show that both the GMC and JB875’s melibiase gene arose via HGT. Our findings enhance the understanding of how metabolic trait variation is generated and maintained.

## Results

### Ethanol as an inducer and the core genes enabling galactose utilization in the *S. pombe* reference background

Consistent with the reported inability of *S. pombe* to utilize galactose, a heterothallic reference background strain failed to grow in yeast extract (YE)-based liquid medium containing galactose as the sole carbon source (*SI Appendix*, Fig. S1*A*). Notably, the utilization of another carbon source, glycerol, by *S. pombe* is stimulated by the supplementation of ethanol (36,37). This prompted us to test whether ethanol could similarly stimulate galactose utilization. Indeed, adding 0.5% ethanol to galactose-containing liquid media enabled growth, whereas YE + 0.5% ethanol did not support growth, consistent with prior reports that ethanol cannot serve as a sole carbon source for *S. pombe* (*SI Appendix*, Fig. S1*A*) (38,39). On solid galactose media, weak growth occurred even without ethanol and was markedly enhanced by its addition (Fig. 1*B*). The reason for this difference between solid and liquid media is unclear. Given the clear visual readout and technical simplicity of solid media, we used YE-based solid media for subsequent assays.

**Figure 1.**
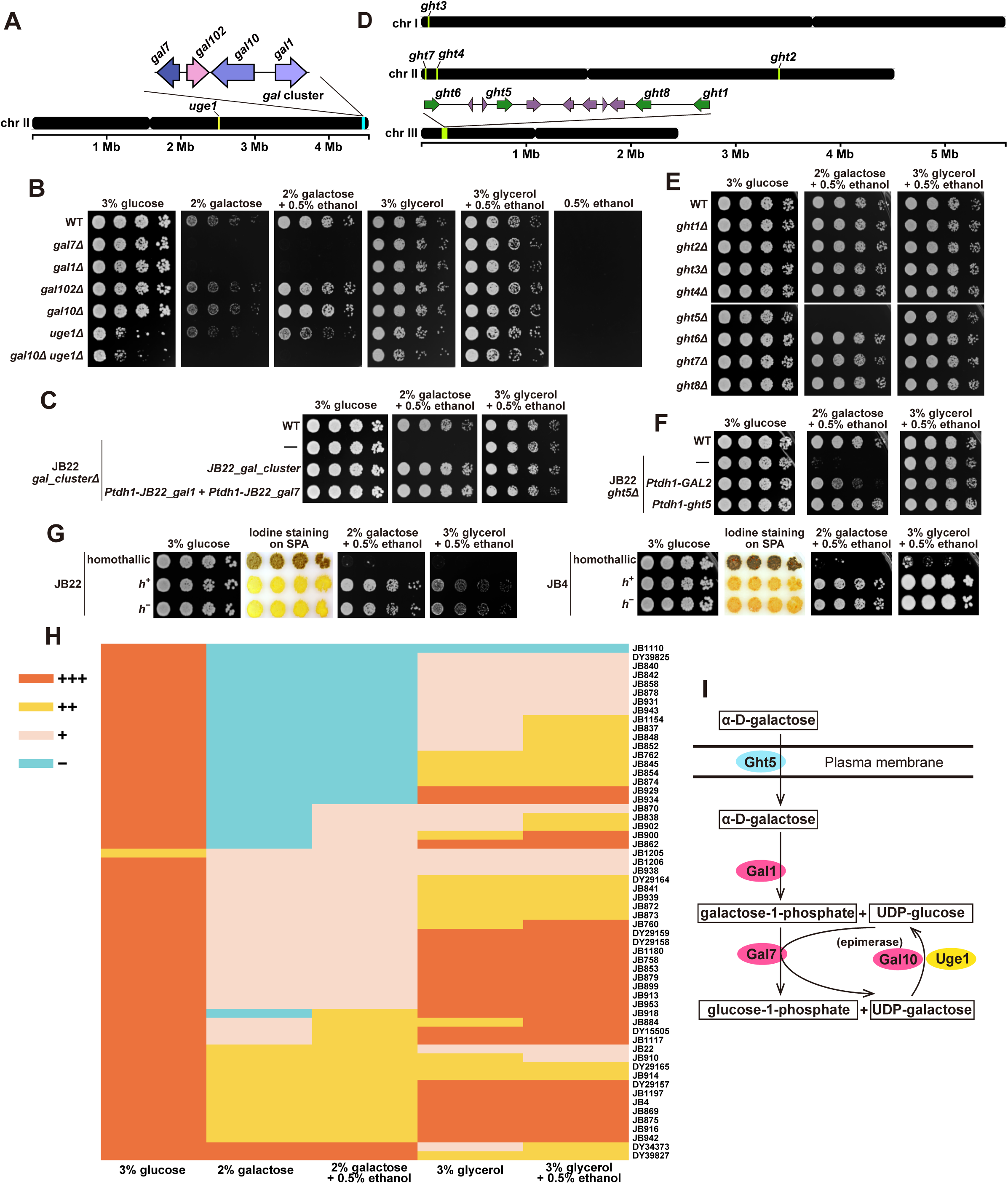
Identification of genes essential for galactose utilization in *S. pombe* reference background and a survey of the galactose utilization trait variation among 58 *S. pombe* natural isolates. (*A*) Locations of the *gal* cluster and the *uge1* gene in the *S. pombe* reference genome. A zoom-in view of the *gal* cluster is shown. (*B*) In the *S. pombe* reference background, *gal1* and *gal7* are essential for galactose utilization, whereas *gal10* and *uge1* are redundantly required for galactose utilization. All media shown here and elsewhere are yeast extract (YE)-based, unless otherwise noted. (*C*) The galactose utilization defect caused by the loss of the *gal* cluster can be rescued by co-expressing *gal1* and *gal7*. (*D*) The locations of eight hexose transporter family genes (*ght1-ght8*) in the *S. pombe* reference genome. A zoom-in view of a chromosome III region containing four *ght* genes is shown, with *ght* genes colored green and non-*ght* genes colored purple. (*E*) *ght5* is required for galactose utilization in the *S. pombe* reference background. (*F*) Expressing Ght5 from the strong promoter *Ptdh1* fully rescued the galactose utilization defect of *ght5Δ*, while expressing the budding yeast galactose transporter Gal2 from the same promoter partially rescued the defect. (*G*) In the reference (JB22) and JB4 backgrounds, homothallic strains exhibited minimal growth on media containing galactose or glycerol as the only carbon source, whereas heterothallic strains exhibited robust growth on the same media. Iodine staining of cells on the standard nitrogen-free sporulation medium SPA showed that homothallic strains are capable of self-mating and sporulation, while heterothallic strains lack such capabilities. Iodine stains starch-like materials in the spore wall. (*H*) A heatmap showing the growth phenotypes of 58 distinct natural isolates of *S. pombe* on five different YE-based solid growth media. Strains used in this analysis are either naturally sterile (JB1110, JB852, JB762, JB929, JB838, JB1206, JB758, DY15505, JB914, and JB869) or are heterothallic derivatives of homothallic isolates. Based on visual inspection of colony growth, the growth phenotype was classified as no obvious growth (–), weak growth (+), moderate growth (++), or robust growth (+++). (*I*) A diagram depicting the galactose utilization pathway in *S. pombe*.

The *gal* cluster on chromosome II contains four genes: *gal7*, *gal102*, *gal10*, and *gal1* (Fig. 1*A*). *S. pombe* Gal10, like its *Sac. cerevisiae* ortholog, harbors both galactose mutarotase and UDP-galactose 4-epimerase domains (9,40,41). Notably, unlike *Sac. cerevisiae*, *S. pombe* possesses a second epimerase gene, *uge1*, which lacks the mutarotase domain and is located far away from the *gal* cluster (Fig. 1*A*) (41). To define the genetic requirement for galactose utilization, we deleted each of these five genes individually. Only deletion of *gal7* or *gal1* abolished growth on galactose (Fig. 1*B*). As expected from the requirement of epimerase activity for galactose utilization, simultaneous deletion of both *gal10* and *uge1* also eliminated growth (Fig. 1*B*). Furthermore, the galactose utilization defect in a *gal* cluster-deleted (*gal_clusterΔ*) strain was fully rescued by ectopically integrating two separate plasmids: one containing *gal1* and another containing *gal7* (Fig. 1*C*). These results demonstrate that, as in *Sac. cerevisiae* (13–15), galactose utilization in *S. pombe* requires Gal1, Gal7, and UDP-galactose 4-epimerase activity—although in *S. pombe*, this activity is redundantly provided by Gal10 or Uge1 (Fig. 1*I*).

*Sac. cerevisiae* uses Gal2, which belongs to the hexose transporter family, for galactose uptake (41). The *S. pombe* reference genome encodes eight hexose transporter family genes (*ght1-ght8*) (Fig. 1*D*) (42). Individual deletions revealed that only *ght5Δ* abolished the galactose utilization ability (Fig. 1*E*). Reintroducing *ght5* or overexpressing *ght1* completely rescued *ght5Δ*, while overexpressing *ght6* conferred partial rescue; no other *ght* genes rescued *ght5Δ* when overexpressed (Fig. 1*F* and *SI Appendix*, Fig. S1*B*). Thus, Ght5 is the essential hexose transporter for galactose utilization in the reference background (Fig. 1*I*), while Ght1 and Ght6 can support galactose utilization only when overexpressed. Notably, *ght1*, *ght5*, and *ght6* are clustered on the left arm of chromosome III (Fig. 1*D*).

### Systematic analysis of galactose utilization variation across natural isolates of *S. pombe*

We next assessed intraspecific variation in galactose utilization using 58 natural isolates: 49 strains representing non-clonal isolates defined by Jeffares et al. (JB strains) (24), and 9 additional distinct isolates (DY strains) (21). These largely cover the known genetic diversity of *S. pombe* (21,24). Within the common callable genomic regions shared by all strains, the pairwise single-nucleotide polymorphism (SNP) distances range from 1,607 to 53,134 (0.014% to 0.465%).

Natural isolates of *S. pombe* are typically homothallic (capable of mating-type switching) (26). By analyzing strains belonging to two distinct backgrounds, JB22 (reference background) and JB4, we found that homothallic strains of both backgrounds efficiently underwent sexual reproduction and sporulation not only on standard nitrogen-free sporulation medium (SPA) (Fig. 1*G* and *SI Appendix*, Fig. S1*C*), but also on YE-based media containing galactose or glycerol as the sole carbon source (*SI Appendix*, Fig. S1*C*). Minimal growth was observed on galactose- and glycerol-containing plates, presumably due to extensive sporulation (Fig. 1*G* and *SI Appendix*, Figs. S1*D* and S3). In contrast, heterothallic strains of these backgrounds exhibited markedly more robust growth on both types of media (Fig. 1 *G* and *H* and *SI Appendix*, Figs. S2 and S3). These results indicate that the propensity of homothallic *S. pombe* strains to undergo sexual reproduction on galactose- or glycerol-containing media interferes with growth-based assays of carbon source utilization. Among the 58 isolates, 10 are naturally heterothallic or sterile (*SI Appendix*, Dataset S1). To ensure consistent and reliable phenotyping, we converted the remaining 48, which are homothallic, to heterothallic. Of these, 31 showed enhanced growth on galactose or glycerol after conversion (Fig. 1*H* and *SI Appendix*, Figs. S1*D* and S3). Growth phenotypes of *h^+^* and *h^−^* heterothallic derivatives were identical (*SI Appendix*, Fig. S3). For these 48 isolates, their heterothallic derivatives were used for the assessment of galactose and glycerol utilization abilities.

One isolate (JB1110) failed to grow on either galactose or glycerol, suggesting a general defect in utilizing non-preferred carbon sources. Of the other 57 isolates—all capable of utilizing glycerol—17 showed no growth on galactose, even with ethanol supplementation. We classified these as Gal^−^. Gal^−^ isolates were scattered across an SNP-based phylogenetic tree, suggesting multiple independent origins of the Gal^−^ trait (*SI Appendix*, Fig. S2). Among galactose-utilizing isolates, two exhibited markedly faster growth than others and were designated Gal^F^. These two Gal^F^ isolates are each other’s closest relatives in the SNP tree (*SI Appendix*, Fig. S2). Together, these results reveal that galactose utilization is a highly variable trait among *S. pombe* natural isolates.

### Seven Gal^−^ isolates harbor naturally occurring *gal* gene deletions

To determine the genetic basis of the Gal^−^ phenotype, we analyzed genomic variation in the *gal* cluster region across the 58 natural isolates (Fig. 2*A*). Seven isolates carried partial or complete deletions of the *gal* cluster—all exhibiting the Gal^−^ phenotype—indicating that gene loss is a direct cause of galactose non-utilization. Based on deletion extent, we classified the deletions into three types: type I (JB840), type II (JB874), and type III (JB854, JB845, JB934, JB943, and JB929). Despite identical deletion boundaries, the five type III isolates carry four distinct SNP signatures in the sequences flanking the deletion (Fig. 2*A*), suggesting at least four independent origins.

**Figure 2.**
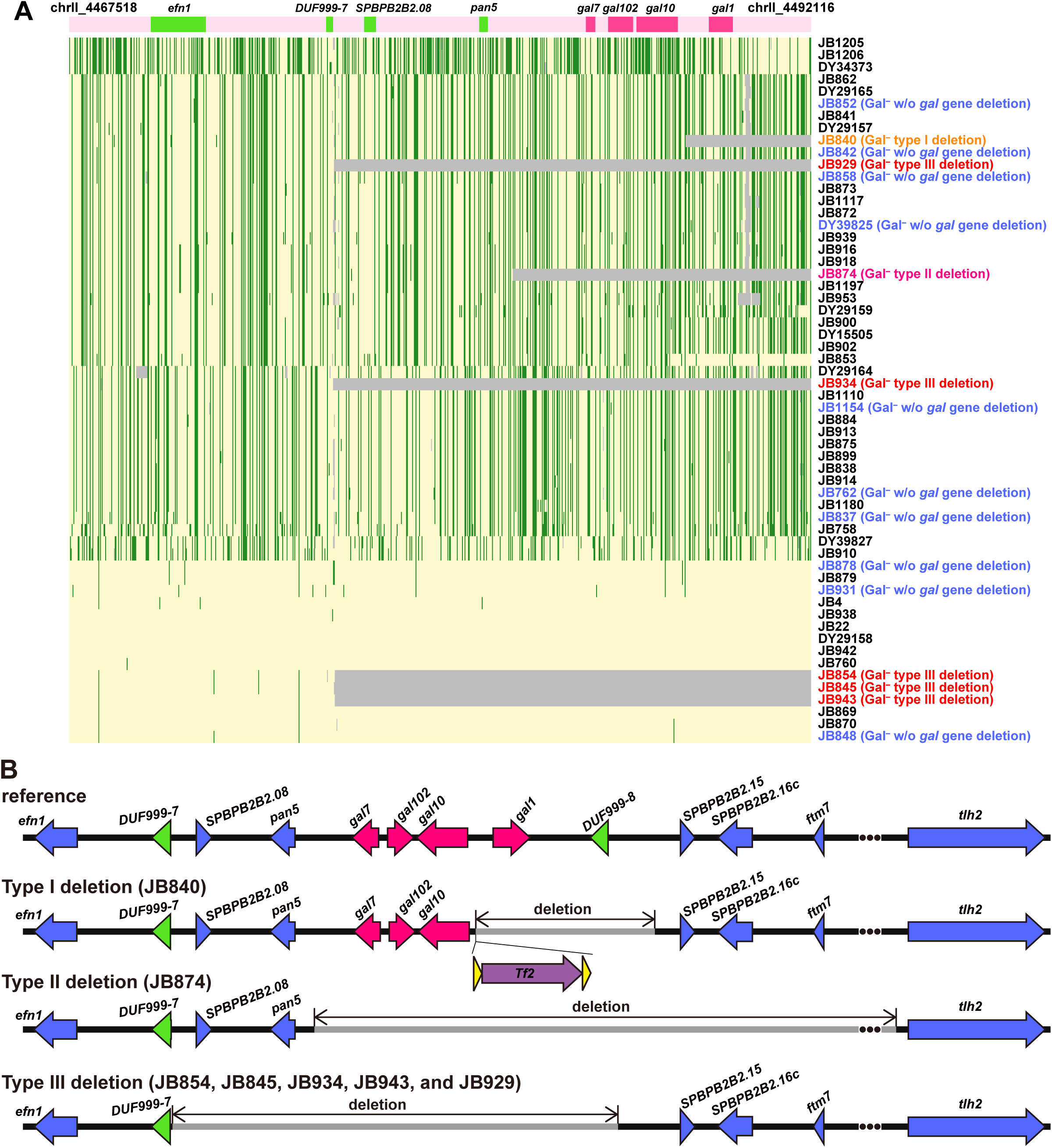
The Gal^−^ trait of seven natural isolates can be attributed to naturally occurring *gal* gene deletions. (*A*) A heatmap showing the SNP patterns in a chromosome II region containing the *gal* cluster in 58 *S. pombe* natural isolates. Each column represents a bi-allelic SNP site, with light yellow indicating the reference allele, green indicating the variant allele, and grey indicating missing values. Genes in this region are shown above the heatmap. The right boundary of this region is immediately adjacent to the *SH* (subtelomeric homologous) region, where SNPs cannot be reliably called due to the repetitive nature of the sequence. (*B*) Schematic diagrams showing the three different types of naturally occurring *gal* gene deletions. The purple arrow represents the internal sequence of a *Tf2* retrotransposon. The yellow triangles represent LTRs of the *Tf2* retrotransposon. Green triangles represent *DUF999* genes (the systematic ID of *DUF999-7* in PomBase is *SPBPB2B2.07c*, and the systematic ID of *DUF999-8* is *SPBPB2B2.14c)*. The *DUF999* family is annotated by PomBase as a *S. pombe*-specific gene family with no known function. Genes in the *gal* cluster are shown as pink arrows.

We characterized deletion junctions to infer deletion mechanisms (Fig. 2*B* and *SI Appendix*, Fig. S4). In type I, *gal1* and the adjacent gene *DUF999-8* (*SPBPB2B2.14c*) are replaced by a full-length Tf2 retrotransposon (Fig. 2*B*). Notably, no target site duplications (TSDs) were detected flanking the retrotransposon (*SI Appendix*, Fig. S4*A*), suggesting that the deletion arose via recombination between two pre-existing Tf2 elements (*SI Appendix*, Fig. S4*B*).

Type II deletes the entire *gal* cluster and all intervening genes between the cluster and *tlh2*, the gene closest to the telomere (Fig. 2*B*). The deletion junction contains a 3-bp microhomology (TTT) (*SI Appendix*, Fig. S4*C*), which may have facilitated deletion formation (*SI Appendix*, Fig. S4*D*).

Type III deletes the entire *gal* cluster and three neighboring genes (Fig. 2*B*). The deletion breakpoints map to highly homologous sequences within two *DUF999* family genes—*DUF999-7* and *DUF999-8*—located on opposite sides of the cluster and oriented in the same direction (Fig. 2*B* and *SI Appendix*, Fig. S4 *E* and *F*). Homologous recombination between these two *DUF999* genes likely caused this recurrent deletion (*SI Appendix*, Fig. S4*G*).

### Ten Gal^−^ isolates possess an intact *gal* cluster but fail to express *gal* genes adequately

For the remaining ten Gal^−^ isolates with an intact *gal* cluster, introducing each of their clusters into a reference background (JB22)-derived *gal_clusterΔ* strain fully restored galactose utilization (Fig. 3 *E* and *H* and *SI Appendix*, Fig. S5 *A-H*), confirming that these clusters are functionally intact.

**Figure 3.**
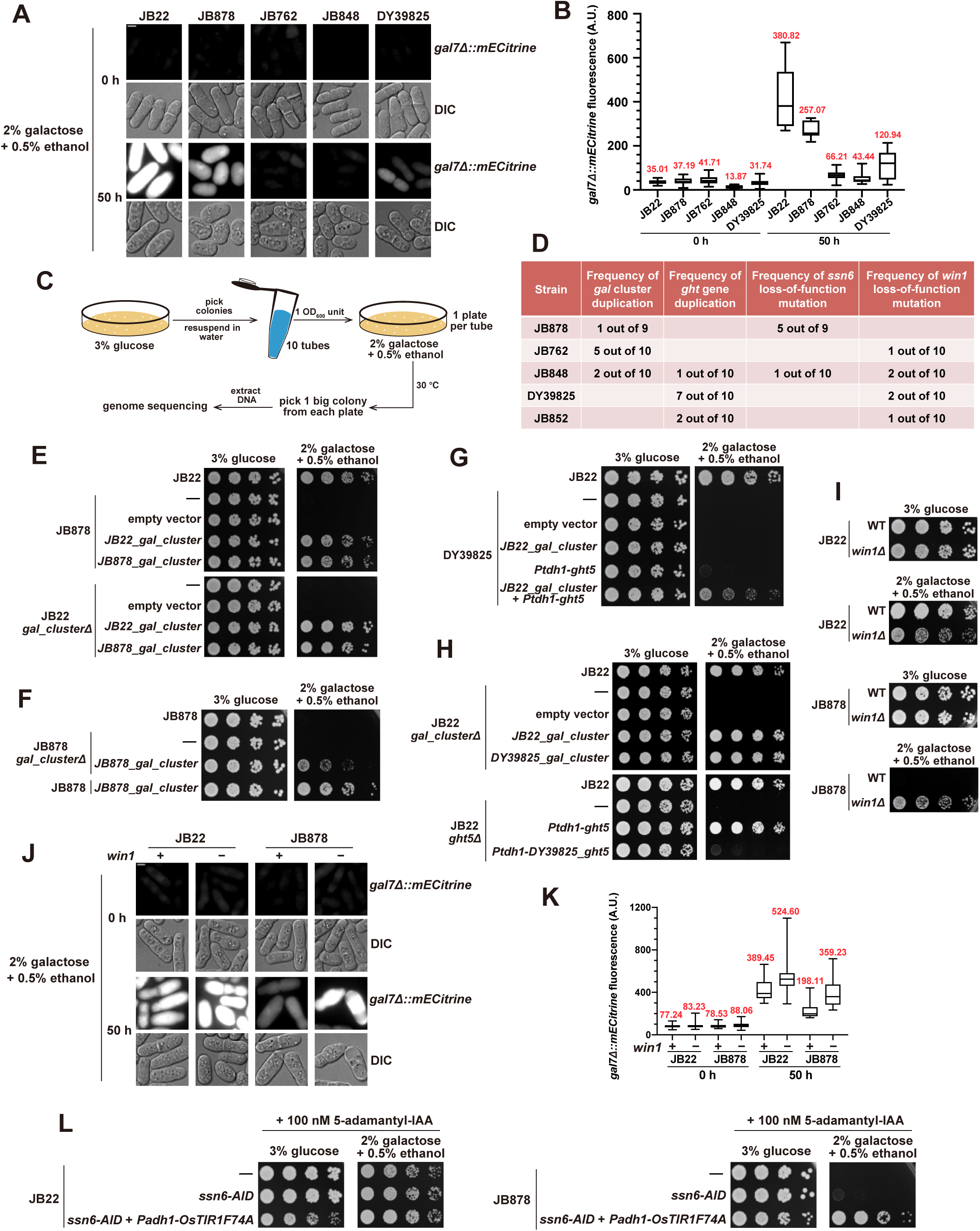
The Gal^−^ trait of 10 natural isolates with an intact *gal* cluster is primarily due to insufficient expression of *gal* genes. (*A*) The levels of *gal7* promoter-driven expression were assessed using an mECitrine reporter integrated at the *gal7* locus, replacing the coding sequence of *gal7* in four Gal^−^ isolates (JB762, JB848, JB878, DY39825) and the reference strain JB22. Scale bar, 3 µm. (*B*) Quantification of mECitrine reporter signals based on imaging data from the experiment shown in (*A*). Median values are shown in red. A.U., arbitrary units. (*C*) Workflow of the spontaneous Gal^+^ revertant screen (see Methods for details). Multiple Gal^−^ background strains were used: one strain with a natural *gal* cluster deletion (JB874), one engineered *gal* cluster deletion (*gal_clusterΔ*) strain in the JB22 background, and five Gal^−^ strains with intact *gal* clusters (JB878, JB762, JB848, DY39825, and JB852). (*D*) Recurrent gene duplications and loss-of-function mutations in spontaneous Gal^+^ mutants. (*E*) JB878 was converted from Gal^−^ to Gal^+^ through introducing an integrating plasmid carrying the *gal* cluster from JB878 (*JB878_gal_cluster*). (*F*) Integrating a plasmid containing *JB878_gal_cluster* into a JB878 strain lacking the endogenous *gal* cluster resulted in a weak Gal^+^ phenotype. (*G*) DY39825 was converted from Gal^−^ to Gal^+^ only when both the *gal* cluster and *ght5* from JB22 were introduced. (*H*) The *gal* cluster but not the *ght5* gene in DY39825 is functional. The *ght5* alleles from DY39825 and JB22 were expressed from the strong *Ptdh1* promoter. (*I*) *win1* deletion weakened the galactose utilization ability of JB22 but converted JB878 from Gal^−^ to Gal^+^. (*J*) *win1* deletion increased the expression of the *gal7Δ::mECitrine* reporter. Scale bar, 3 µm. (*K*) Quantification of mECitrine reporter signals based on imaging data from the experiment shown in (*J*). Median values are shown in red. (*L*) AID (auxin-inducible degron)-based down-regulation of Ssn6 converted JB878 from Gal^−^ to Gal^+^.

To test whether expression is impaired, we integrated an mECitrine reporter under the native *gal7* promoter in four of these ten Gal^−^ isolates (JB762, JB848, JB878, DY39825) and JB22. In glucose, all showed negligible fluorescence (Fig. 3 *A* and *B*). Upon galactose shift, JB22 exhibited strong induction; JB762 and JB848 showed minimal response; JB878 and DY39825 displayed intermediate levels—indicating partial derepression (Fig. 3 *A* and *B*). These results suggest that insufficient *gal* gene expression underlies the Gal^−^ phenotype.

To identify genetic causes unbiasedly, we performed spontaneous Gal^+^ revertant screens across five Gal^−^ isolates with intact clusters (JB878, JB762, JB848, DY39825, JB852) and two control strains lacking the *gal* cluster (JB874 and JB22 *gal_clusterΔ*) (Fig. 3*C*). Revertants arose only in the strains with intact clusters. Whole-genome sequencing of revertants revealed four recurrent mutation classes: (1) *gal* cluster duplications, (2) *ght* gene duplications, (3) loss-of-function mutations in the transcriptional repressor gene *ssn6*, and (4) loss-of-function mutations in the mitogen-activated protein kinase (MAPK) pathway gene *win1* (Fig. 3*D* and *SI Appendix*, Dataset S3).

The frequent *gal* cluster duplications in Gal^+^ revertants suggest that *gal* cluster gene expression is limiting. Introducing into JB878 an integration-plasmid-borne *gal* cluster from JB878 restored growth (Fig. 3*E*), but growth was weaker when the same plasmid was introduced into a JB878 *gal_clusterΔ* strain (Fig. 3*F*). This indicates that both endogenous and ectopic copies contribute to full galactose utilization—implying that the endogenous genes are expressed, but below a functional threshold. The plasmid performs better in JB22 *gal_clusterΔ* than in JB878 *gal_clusterΔ* (Fig. 3 *E* and *F*), suggesting that trans-acting factors in JB878 limit expression—consistent with recurrent *ssn6* mutations in its revertants (Fig. 3*D* and *SI Appendix*, Dataset S3).

Introducing an additional *gal* cluster copy converted 8 of the 10 Gal^−^ isolates to Gal^+^ (Fig. 3*E* and *SI Appendix*, Fig. S5 *A-G*), indicating that insufficient expression of the *gal* genes causes the Gal^−^ phenotype. The two exceptions—DY39825 and JB852—remained Gal^−^ even after *gal* cluster duplication (Fig. 3*G* and *SI Appendix*, Fig. S5*I*), suggesting the existence of other underlying causes. In Gal^+^ revertants of these isolates, we observed duplications of multiple *ght* genes (Fig. 3*D* and *SI Appendix*, Dataset S3), implicating hexose transporters in their phenotype. Indeed, when expressed from the strong *Ptdh1* promoter in a *ght5Δ* JB22 strain, the *ght5* alleles from DY39825 and JB852 failed to restore galactose utilization (Fig. 3*H* and *SI Appendix*, Fig. S5*H*).

To investigate the molecular basis of *ght5* dysfunction, we constructed a phylogenetic tree of *ght* family genes across the 58 isolates (*SI Appendix*, Fig. S6). Strikingly, the *ght5* allele of DY39825 branches immediately adjacent to the *ght8* clade, suggesting recombination between *ght5* and *ght8*. Amino acid alignment revealed that positions 335 and 427—alanine and asparagine in all functional *ght5* alleles—are replaced by glycine and aspartate in DY39825, matching the *ght8* sequence (*SI Appendix*, Fig. S7). These substitutions may render the transporter unable to import galactose. The basis of *ght5* dysfunction in JB852 remains unclear.

Despite this defect, expressing the functional *ght5* allele from JB22 only weakly improved galactose utilization in DY39825 and failed to rescue JB852 (Fig. 3*G* and *SI Appendix*, Fig. S5*I*), suggesting additional limitations. We hypothesized that these strains also suffer from suboptimal *gal* gene expression. Indeed, introducing both the JB22 *gal* cluster and *ght5* gene further enhanced galactose utilization in DY39825 and allowed weak growth of JB852 on galactose (Fig. 3*G* and *SI Appendix*, Fig. S5*I*). Thus, the Gal^−^ phenotype in DY39825 and JB852 results from combined defects in galactose transport and *gal* gene expression.

*win1* encodes a MAPK kinase kinase (MAPKKK) upstream of the stress-responsive MAPK Sty1 (43). We found that *win1Δ* converted multiple Gal^−^ isolates to Gal^+^ (Fig. 3*I* and *SI Appendix*, Fig. S5*J*), yet reduced galactose utilization in JB22 (Fig. 3*I*), revealing a strain-background-dependent effect. Using the *gal7Δ::mECitrine* reporter, we found that *win1Δ* increased *gal7* expression in JB878 (Fig. 3 *J* and *K*), suggesting that Win1 represses the *gal* genes in certain backgrounds. *sty1Δ* also improved galactose utilization in Gal^−^ isolates, albeit less robustly (*SI Appendix*, Fig. S5 *J* and *K*), consistent with its single isolation in the screen.

Premature stop codons in *ssn6* (occurring no earlier than codon 621) were isolated five times in JB878 and once in JB848 (Fig. 3*D* and *SI Appendix*, Dataset S3). Since *ssn6* is essential, these likely cause partial loss of function. Degron-mediated depletion of Ssn6 converted JB878 to Gal^+^ (Fig. 3*L*), directly implicating this transcriptional repressor in the Gal^−^ phenotype.

### A tandemly repeated, lineage-specific gene cluster drives the Gal^F^ phenotype

Among 58 isolates, DY39827 and DY34373 exhibited exceptionally fast growth on galactose and were designated Gal^F^ (Figs. 1*H* and 4*A* and *SI Appendix*, Figs. S2 and S3). Deletion of the canonical *gal* cluster did not impair galactose utilization in DY39827 (Fig. 4*B*), suggesting that an alternative locus drives the Gal^F^ phenotype.

**Figure 4.**
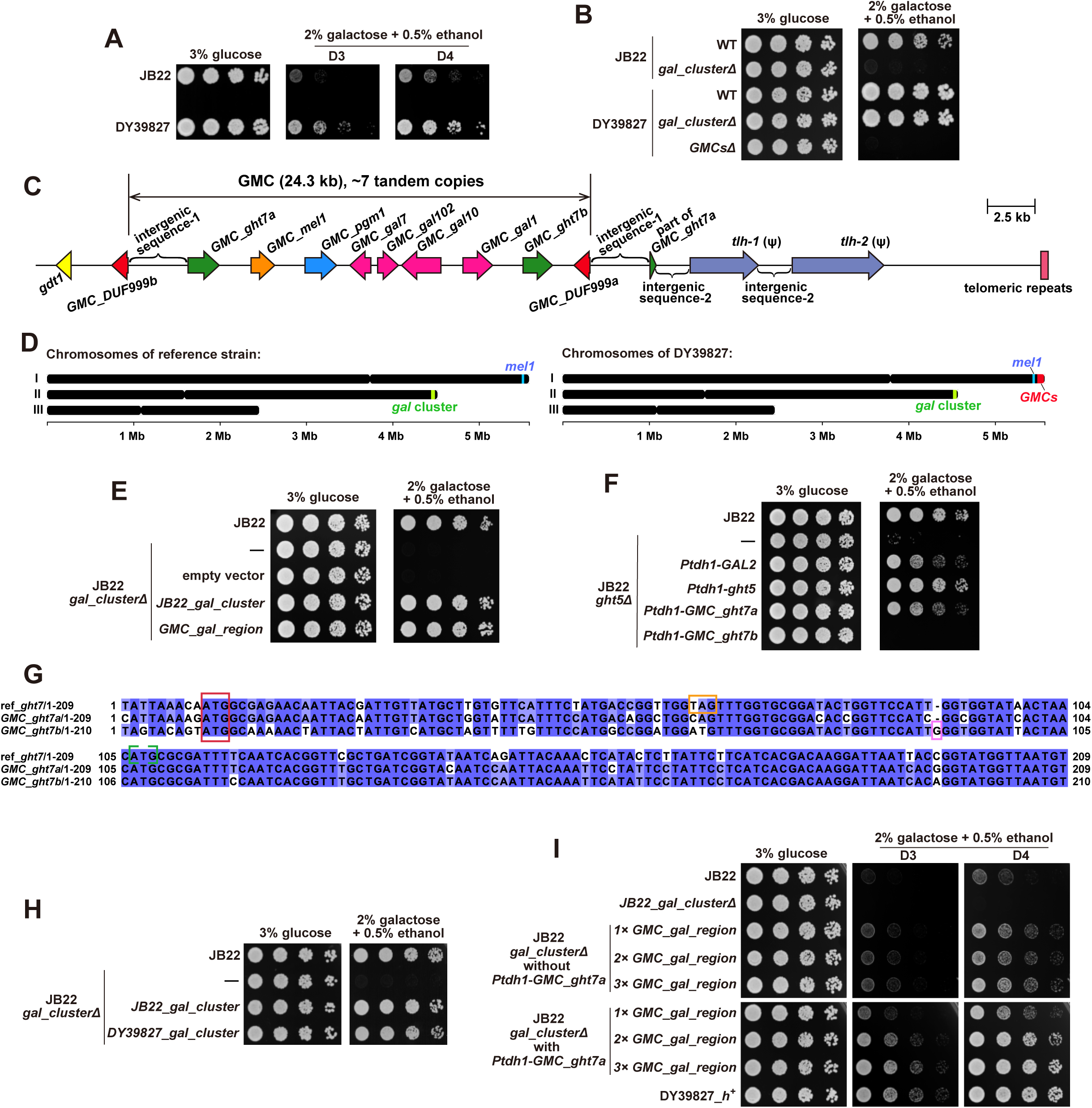
The *gal-mel* clusters (GMCs) in DY39827 underlie the Gal^F^ trait. (*A*) DY39827 exhibited a markedly higher growth rate on galactose media than JB22. Images taken on day 3 (D3) and day 4 (D4) are shown. (*B*) The loss of the GMCs but not the *gal* cluster abolished the galactose utilization ability of DY39827. (*C*) A diagram depicting the chromosome I right-end region containing the GMCs in DY39827. The symbol “Ψ” indicates a pseudogene. The DY39827 genome was sequenced using PacBio HiFi technology. (*D*) Diagrams showing the genomic locations of the *gal* cluster, the *mel1* gene, and the GMCs. (*E*) The *GMC_gal_region* can rescue the galactose utilization defect of a JB22 strain lacking the *gal* cluster. The *GMC_gal_region* is a GMC segment that contains the four *gal*-homologs. It was cloned into an integrating plasmid and introduced into a *gal_clusterΔ* JB22 strain. (*F*) *GMC_ght7a* but not *GMC_ght7b* can rescue the galactose utilization defect of a JB22 strain lacking *ght5*. (*G*) Sequence alignment of *ght7* in the reference genome (ref*_ght7*), *GMC_ght7a*, and *GMC_ght7b*. The sequences all begin at the 9^th^ base upstream of the start codon. Identical bases are highlighted with a blue background. The red box highlights the start codons, while the orange box denotes an early stop codon at position 19 in ref*_ght7*, rendering it a pseudogene. The green bracket indicates the incorrectly annotated start codon of ref_*ght7* in PomBase. The light purple box shows the frameshifting one-base insertion in *GMC_ght7b* (insertion of ‘G’ after codon 27). (*H*) The *DY39827_gal_cluster* (the canonical chromosome II *gal* cluster in DY39827) can rescue the galactose utilization defect of a JB22 strain lacking the *gal* cluster. (*I*) Combining multiple copies of *GMC_gal_region* with *Ptdh1*-driven *GMC_ght7a* enabled JB22 to grow as well as DY39827 on galactose media.

We generated a PacBio HiFi genome assembly of DY39827. It revealed a tandem repeat array near the right telomere of chromosome I, containing nine genes per 24.3-kb repeat unit (Fig. 4 *C* and *D*). The nine genes include homologs similar but not identical to each of the four genes in the canonical *gal* cluster, two *ght7*-homologs, a *mel1*-homolog (encoding melibiase that hydrolyzes melibiose), a *pgm1*-homolog (encoding phosphoglucomutase acting downstream of the Leloir enzymes), and a *DUF999*-family gene (Fig. 4*C*). Given that most of the nine genes are related to galactose/melibiose utilization, we named this repeat unit the *gal-mel* cluster (GMC). The order of genes is: *GMC_ght7a-GMC_mel1-GMC_pgm1-GMC_gal7-GMC_gal102-GMC_gal10-GMC_gal1-GMC_ght7b-GMC_DUF999a*.

Notably, an additional *DUF999* gene—*GMC_DUF999b*—with 97% nucleotide identity (99.41% excluding a 21-bp gap) to *GMC_DUF999a* (*SI Appendix*, Fig. S8*A*), lies immediately on the centromere side of the first GMC copy (Fig. 4*C*). Thus, each GMC copy is flanked by identical or highly similar *DUF999* genes, potentially facilitating recombination and copy number variation.

BLAST analysis against Illumina genome assemblies of the 58 isolates identified the GMC in four isolates: the two Gal^F^ isolates DY39827 and DY34373, and two other isolates, JB1205 and JB1206—three of which originated from the former Soviet Union and the other from Ukraine (*SI Appendix*, Dataset S1). Quantitative PCR showed that the GMC is present in 4 to 8 copies in these isolates (Fig. 4*C* and *SI Appendix*, Fig. S8 *B* and *C*). Deletion of all GMC copies in DY39827 abolished galactose utilization (Fig. 4*B*), demonstrating that the GMC is essential for galactose utilization in this background.

To test functional equivalence, we introduced the *GMC_gal_region* (containing the four *gal*-homologs) into a *gal_clusterΔ* JB22 strain. It fully restored galactose utilization, matching the canonical *JB22_gal_cluster* (Fig. 4*E*). Of the two *ght7* homologs, only *GMC_ght7a* restored growth in a *ght5Δ* JB22 strain on galactose (Fig. 4*F*). Sequence analysis revealed that *GMC_ght7b* and the reference *ght7* are pseudogenes (Fig. 4*G*). Even though the canonical *gal* cluster in DY39827 does not support galactose utilization in its native background (Fig. 4*B*), when introduced into *gal_clusterΔ* JB22, it rescued galactose utilization (Fig. 4*H*), indicating repression by trans-acting factors.

Introducing two or three copies of *GMC_gal_region* into *gal_clusterΔ* JB22 did not result in stronger growth on galactose than introducing a single copy (Fig. 4*I*). Co-expression of *GMC_ght7a* increased the growth rate proportionally with the copy number of *GMC_gal_region*. With three copies of *GMC_gal_region* plus *GMC_ght7a*, growth matched DY39827 (Fig. 4*I*). These findings demonstrate that high gene dosage due to the multicopy nature of the GMC drives the Gal^F^ phenotype.

### Lineage-specific melibiase genes enable melibiose utilization in a minority of *S. pombe* isolates

The GMC also contains *GMC_mel1*, encoding a melibiase that hydrolyzes melibiose into galactose and glucose (Figs. 4*C* and 5*A*). *GMC_mel1* is only distantly related to the canonical *mel1* gene present in all 58 isolates, sharing 71% nucleotide identity with the canonical *mel1* in the reference genome (*JB22_mel1*) (Fig. 4*D*). We systematically assayed melibiose utilization across the 58 isolates. Only five exhibited robust growth on melibiose (Mel^+^)—including all four GMC-containing isolates and another isolate, JB875, which originated from Spain (Fig. 5*B* and *SI Appendix*, Fig. S9*A* and Dataset S1).

**Figure 5.**
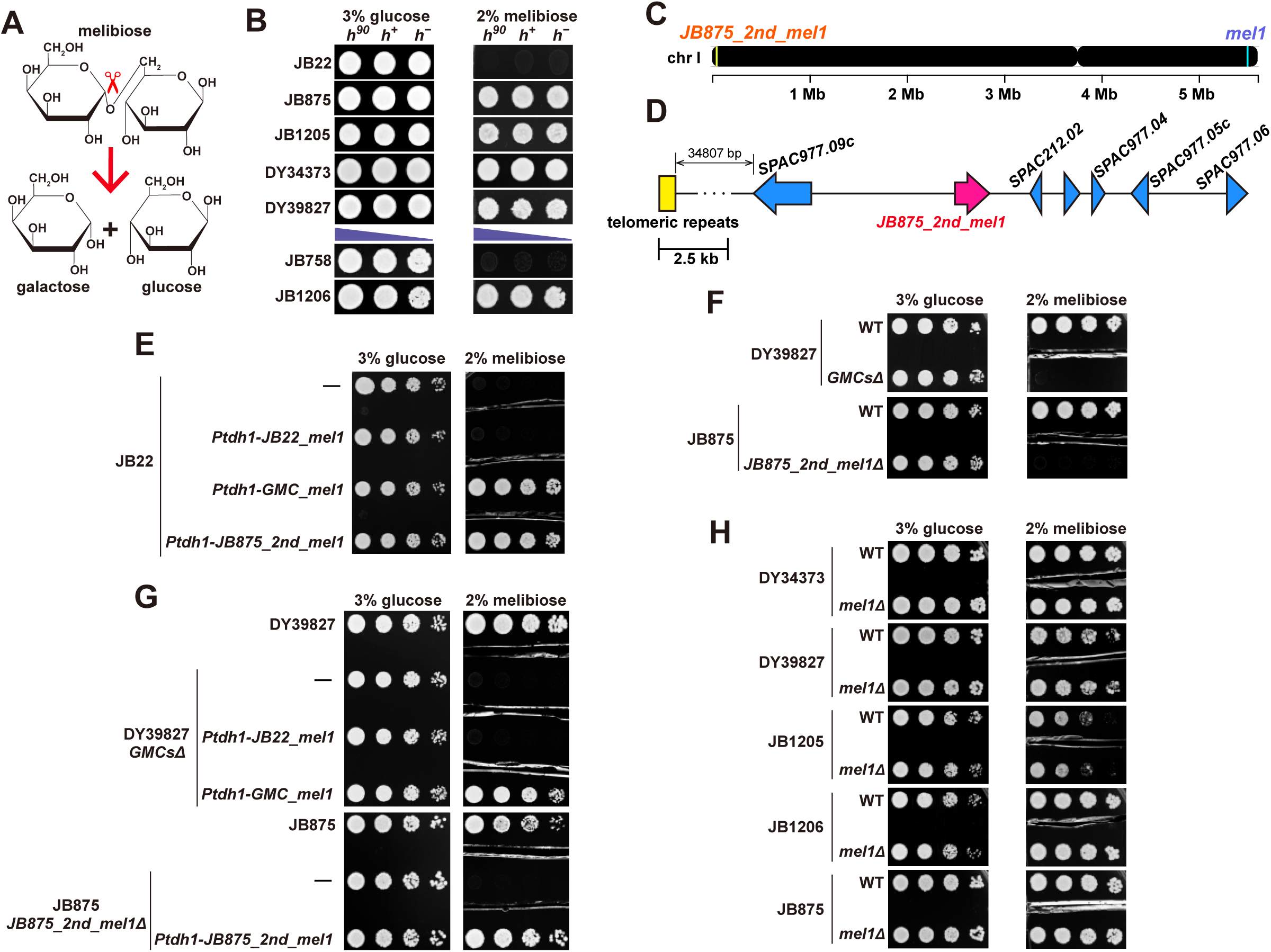
Functional characterization of two melibiase genes enabling melibiose utilization. (*A*) Schematic representation of the melibiase-catalyzed hydrolysis reaction that converts melibiose into galactose and glucose. (*B*) Only the four strains harboring the GMC (DY34373, DY39827, JB1205, and JB1206) and one additional strain, JB875, can efficiently utilize melibiose as a carbon source (Mel^+^ trait). JB22 and JB758 are shown as representatives of the remaining 53 strains that cannot efficiently utilize melibiose (Mel^−^ trait). For natural isolates that are homothallic (denoted as *h^90^* for brevity), the engineered heterothallic derivatives (*h^+^* and *h^−^*) were also examined. For natural isolates that cannot undergo sexual reproduction (including JB758 and JB1206), multiple dilutions indicated by purple triangles were spotted on the plates. (*C*) A diagram depicting the locations of the *JB875_2nd_mel1* gene and the *mel1* gene on chromosome I of JB875. The JB875 genome was sequenced using PacBio HiFi technology. (*D*) A diagram depicting the *JB875_2nd_mel1* gene and its surrounding region. (*E*) The expression of either *GMC_mel1* or *JB875_2nd_mel1* under the control of the strong promoter *Ptdh1* converted JB22 from Mel^−^ to Mel^+^. The *JB22_mel1* corresponds to the canonical *mel1* gene found in the reference genome. To prevent interference in phenotype determination from the diffusion of extracellular glucose generated by the secreted melibiase, gaps were created on the agar plates by cutting out agar slices. (*F*) The loss of GMCs in DY39827 and the loss of *JB875_2nd_mel1* in JB 875 converted them from Mel^+^ to Mel^−^. (*G*) *GMC_mel1* or *JB875_2nd_mel1*, but not *JB22_mel1*, expressed from *Ptdh1*, restored efficient melibiose utilization in the DY39827 strain lacking the GMCs or JB875 strain lacking the *JB875_2nd_mel1*. (*H*) The loss of the *mel1* gene had no effect on the melibiose utilization abilities of the five Mel^+^ strains.

The PacBio HiFi assembly of JB875 revealed a non-canonical melibiase gene located in the subtelomeric region of the left arm of chromosome I (Fig. 5 *C* and *D* and *SI Appendix*, Fig. S9*C*). This gene shares 70% and 83% nucleotide identity with *JB22_mel1* and *GMC_mel1*, respectively. We designate it *JB875_2nd_mel1*. Illumina read mapping revealed that it is present as a single copy (*SI Appendix*, Fig. S9*B*).

Expression of *GMC_mel1* or *JB875_2nd_mel1* but not *JB22_mel1* from the strong promoter *Ptdh1* conferred robust melibiose utilization in JB22 (Fig. 5*E*). Deletion of the GMC array in DY39827 or *JB875_2nd_mel1* in JB875 abolished melibiose utilization (Fig. 5*F*). Reintroduction of *Ptdh1*-driven *GMC_mel1* into a GMC-deleted DY39827 strain, or *JB875_2nd_mel1* into JB875 strain lacking the *JB875_2nd_mel1* restored the Mel^+^ phenotype (Fig. 5*G*). Notably, deletion of the canonical *mel1* had no effect on Mel^+^ strains (Fig. 5*H*).

Thus, the ubiquitously present canonical *mel1* is non-functional for melibiose catabolism. Melibiose utilization is conferred exclusively by two divergent, lineage-specific melibiase genes: *GMC_mel1* (in four isolates) and *JB875_2nd_mel1* (in one isolate).

### Phylogenetic analysis supports horizontal acquisition of the GMC and *JB875_2nd_mel1*

To infer the evolutionary origins of the GMC and *JB875_2nd_mel1*, we constructed maximum likelihood phylogenies that include homologs across *Schizosaccharomyces* species (Fig. 6). Five GMC genes—*GMC_gal7*, *GMC_gal10*, *GMC_gal1*, *GMC_gal102*, and *GMC_pgm1*—each form a clade sister to their canonical *S. pombe* homologs, with genetic distances substantially exceeding those between *S. octosporus* and *S. lindneri* genes (Fig. 6 *A-E*). This indicates that these genes did not arise from recent intraspecific duplication but likely originated via HGT from an unsampled *Schizosaccharomyces* lineage whose divergence from *S. pombe* exceeds that between known sister species within the genus.

**Figure 6.**
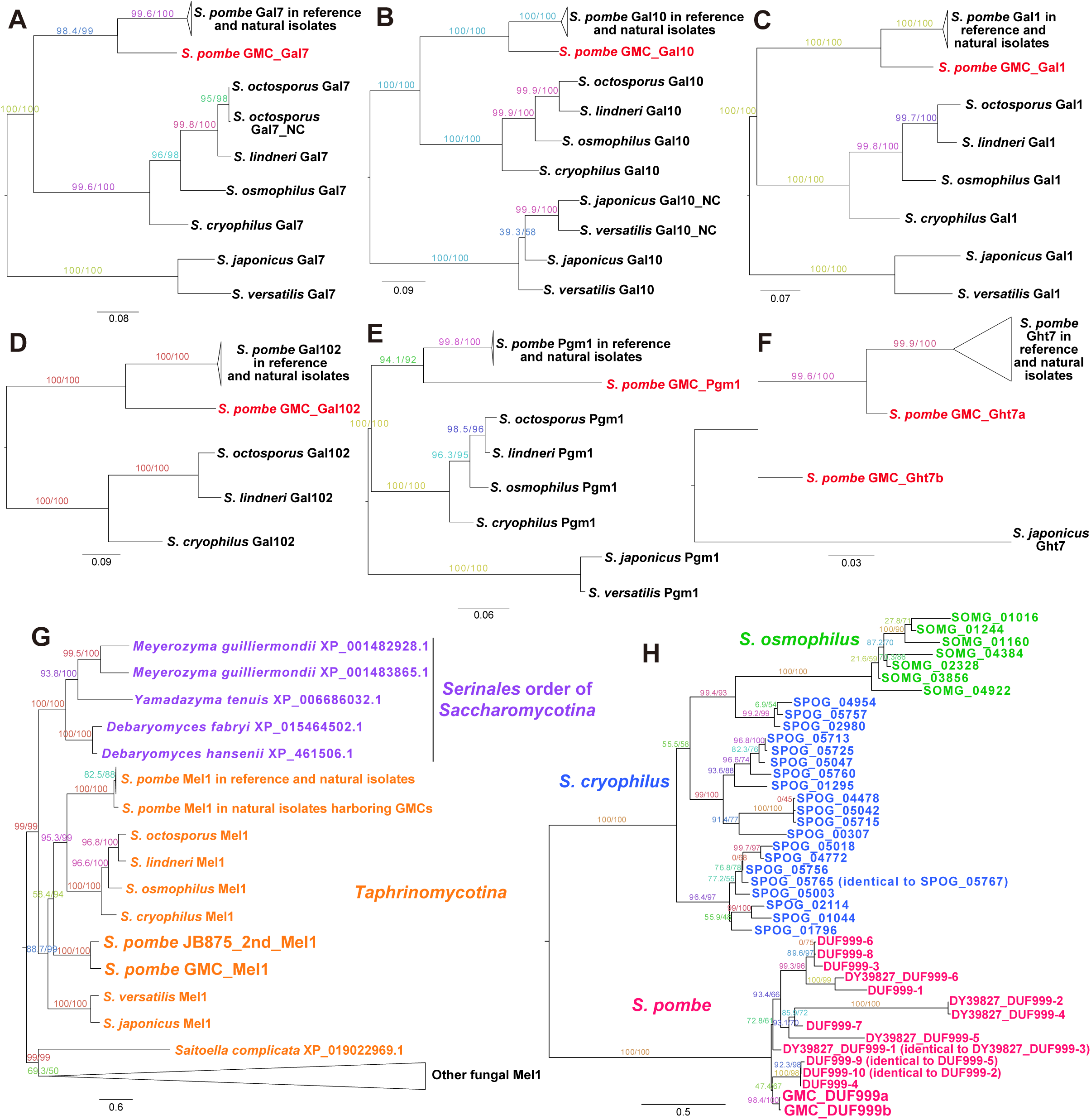
Phylogenetic analyses of GMC genes and the *JB875_2nd_mel1* gene. (*A-H*) Maximum likelihood (ML) phylogenetic trees of GMC_Gal7 and its homologs across fission yeasts (*A*), GMC_Gal10 and its homologs across fission yeasts (*B*), GMC_Gal1 and its homologs across fission yeasts (*C*), GMC_Gal102 and its homologs across fission yeasts (*D*), GMC_Pgm1 and its homologs across fission yeasts (*E*), GMC_Ght7a, GMC_Ght7b, and their closest homologs (*F*), GMC_Mel1, JB875_2nd_Mel1, and their fungal homologs (*G*), and GMC_DUF999a, GMC_DUF999b, and their homologs across fission yeasts (*H*). The trees were constructed using IQ-TREE. All trees, except for the one in (*G*), were rooted using midpoint rooting, while the tree in (*G*) was rooted with melibiases from *S. complicata* and other fungi as the outgroup. Branch support values were calculated by IQ-TREE. The amino acid sequences of homologs of GMC genes and *JB875_2nd_mel1* were obtained through BLAST searches. The appendix “NC” denotes Gal7- and Gal10-coding genes located outside the *gal* cluster. *DUF999* genes in the reference genome and those not associated with the GMCs in the DY39827 genome were numbered according to their chromosomal positions (see *SI Appendix*, Fig. S11).

We observed that the canonical *gal102* gene has undergone multiple independent inactivation or loss events, suggesting relaxation of selective constraint on *gal102*. In three *S. pombe* isolates—DY34373, JB1205, and JB1206—*gal102* contains both an LTR insertion and a frameshift mutation, rendering it a pseudogene (*SI Appendix*, Fig. S10*A*). In *S. osmophilus*, *gal102* is highly degenerate, with numerous inactivating mutations (*SI Appendix*, Fig. S10*B*), while no homolog is detectable in *S. japonicus* or *S. versatilis* (*SI Appendix*, Fig. S10*C*). These observations suggest that *gal102* is under relaxed selective constraint, consistent with our experimental finding that it is dispensable for galactose utilization in *S. pombe*.

*GMC_ght7a* and *GMC_ght7b* are highly divergent from canonical *ght7* genes in *S. pombe* (Fig. 6*F*), but complex paralogy among hexose transporters across fission yeasts precludes a confident assessment of whether their divergence exceeds that typically observed between sister species.

The melibiase phylogeny reveals that *GMC_mel1* and *JB875_2nd_mel1* form a distinct clade despite substantial sequence divergence (Fig. 6*G*). This clade is sister to a monophyletic group containing canonical *mel1* from *S. pombe*, *S. octosporus*, *S. lindneri*, *S. osmophilus*, and *S. cryophilus*. This topology implies that *GMC_mel1* and *JB875_2nd_mel1* derive from a lineage more distantly related to *S. pombe* than the other four species. Notably, the closest homologs of all fission yeast melibiases are found in the budding yeast order *Serinales*, suggesting that ancestral melibiase genes may have undergone HGT across *Ascomycota*. *Serinales* is the donor order for the horizontally transferred *gal* cluster in fission yeasts (5, 9). This convergence suggests that *Serinales* may have served as a recurrent source of metabolic genes for fission yeasts via HGT.

Due to the large number of *DUF999* paralogs, we present a focused phylogeny including representatives from the *S. pombe* reference, the GMC-containing isolate DY39827, and the only other species with identifiable *DUF999* genes: *S. cryophilus* and *S. osmophilus* (Fig. 6*H*). *DUF999* genes are located in subtelomeric regions of chromosomes I and II in both the reference and DY39827 (*SI Appendix*, Fig. S11). In the tree, *GMC_DUF999a* and *GMC_DUF999b* are nested within a clade of *S. pombe DUF999* genes, indicating that, unlike the other GMC genes, they are of *S. pombe* origin (Fig. 6*H*).

### The chromosomal region adjacent to the GMC array exhibits elevated sequence divergence and mosaic evolutionary origins

In the 4 GMC-containing isolates, the tandemly arrayed GMC copies are located near the right telomere of chromosome I. Comparative analysis of genome assemblies revealed that, on the centromere-proximal side of the GMC array, these isolates maintain near-perfect synteny with the reference genome. However, their nucleotide sequences, which are nearly identical to each other, exhibit unusually strong divergence from the reference genome across an ∼120-kb region adjacent to the GMC array, which we designate the “syntenic divergent region” (*SI Appendix*, Fig. S12 *A-C*).

To quantify this divergence at the gene level, we calculated pairwise *ds* (synonymous substitution rate) values for syntenic protein-coding genes in this region. As a control, *ds* values between representatives of the two ancient lineages of *S. pombe*—the reference, representing the REF lineage, and JB864, representing the NONREF lineage—are relatively uniform across this region, averaging ∼0.019. This corresponds to the highest level of intraspecific variation. Strikingly, *ds* values between the reference and the GMC-containing isolate DY39827 are dramatically higher in this region, averaging ∼0.214 (*SI Appendix*, Fig. S12*C*). Notably, divergence is not uniform: a 12-gene subregion (spanning *SPAC29B12.11c* to *SPAC1039.08*) shows exceptionally high *ds* values, averaging ∼0.361, suggesting a mosaic genomic architecture with mixed evolutionary origins. For two conserved genes within the high-*ds* subregion—*SPAC29B12.11c* and *SPAC29B12.12* (*hot13*)—the divergence between the reference and DY39827 reaches that observed between the sister species *S. octosporus* and *S. lindneri* (*SI Appendix*, Fig. S12 *D-F*).

Together, these findings indicate that this divergent chromosomal region adjacent to the GMC array has a complex evolutionary origin. The GMC may have first arrived in a divergent, unsampled *S. pombe* lineage before a co-transfer with the adjacent region via introgression.

### Cross-species synteny analysis reveals dynamic evolution of galactose and melibiose utilization genes

To gain insight into the evolutionary dynamics of galactose and melibiose utilization genes, we performed cross-species synteny analysis across fission yeasts (*SI Appendix*, Fig. S13). The *gal* cluster—comprising *gal7*, *gal102*, *gal10*, and *gal1*—is conserved in gene order and orientation across all species, with the exception of *gal102*, which is absent in *S. japonicus* and *S. versatilis* (*SI Appendix*, Fig. S13*A*). In contrast, the genomic regions flanking the *gal* cluster exhibit three distinct synteny patterns: one shared by *S. octosporus*, *S. lindneri*, *S. osmophilus*, and *S. cryophilus*; a second specific to *S. pombe*; and a third shared by *S. japonicus* and *S. versatilis*. This indicates that the *gal* cluster has undergone at least two independent genomic relocations during fission yeast evolution without disruption of the cluster itself, suggesting strong selective pressure to maintain its integrity as a functional unit.

The four genes in the *GMC_gal_region* share the same gene order and orientation as the canonical *gal* cluster genes. Notably, *GMC_DUF999a* and *GMC_DUF999b* are oriented in the same direction as the two *DUF999* genes located on opposite sides of the canonical *gal* cluster in *S. pombe* (Figs. 2*B* and 4*C* and *SI Appendix*, Fig. S13*A*). This conserved arrangement suggests that the GMC may have originated at the canonical *gal* locus and later moved to its current location through ectopic recombination mediated by *DUF999* genes. Recombination between *DUF999* genes likely also underlies both the tandem amplification of the GMC and the type III deletions of the canonical *gal* cluster observed in Gal^−^ strains (Figs. 2*B* and 4*C* and *SI Appendix*, Fig. S4 *E-G*).

We identified two lineage-specific duplications of individual *gal* genes (Fig. 6 *A* and *B* and *SI Appendix*, Fig. S13*A*). In *S. japonicus* and *S. versatilis*, an additional copy of *gal10*—designated *gal10_NC*—is present outside the canonical cluster. Similarly, *S. octosporus* carries an extra copy of *gal7*, designated *gal7_NC*, located outside the cluster. Intriguingly, a hexose transporter gene is located immediately adjacent to *gal10_NC* in *S. japonicus* and *S. versatilis*, suggesting possible functional linkage or co-regulation.

Similar to the flanking regions of the *gal* cluster, the genomic context of *mel1* also exhibits three distinct synteny patterns corresponding to the same species groupings (*SI Appendix*, Fig. S13*B*). Strikingly, in *S. japonicus* and *S. versatilis*, *mel1* is located directly adjacent to the *gal* cluster, forming a single, larger gene cluster that encompasses both galactose and melibiose utilization genes. This configuration resembles the structure of the GMC, indicating that the physical linkage of these metabolic functions has evolved convergently in distant lineages.

*uge1* and *pgm1*, which not only contribute to galactose utilization but also perform essential cellular functions beyond this role, exhibit more conserved synteny across species (*SI Appendix*, Fig. S13 *C* and *D*). They may be under stronger purifying selection, which could account for their more stable genomic positions compared to the more dynamically evolving *gal* and *mel1* loci.

### Efficient melibiose and galactose utilization confers an advantage in raffinose-rich environments

Raffinose—a trisaccharide composed of galactose, glucose, and fructose—is considered the second most abundant water-soluble carbohydrate in nature after sucrose (44,45). Acting as a reserve carbohydrate and a stress-tolerance factor in plants, it accumulates to high levels in many plant tissues and organs (46,47). Raffinose can be hydrolyzed into melibiose and fructose by invertase or into galactose and sucrose by melibiase (Fig. 7*A*).

**Figure 7.**
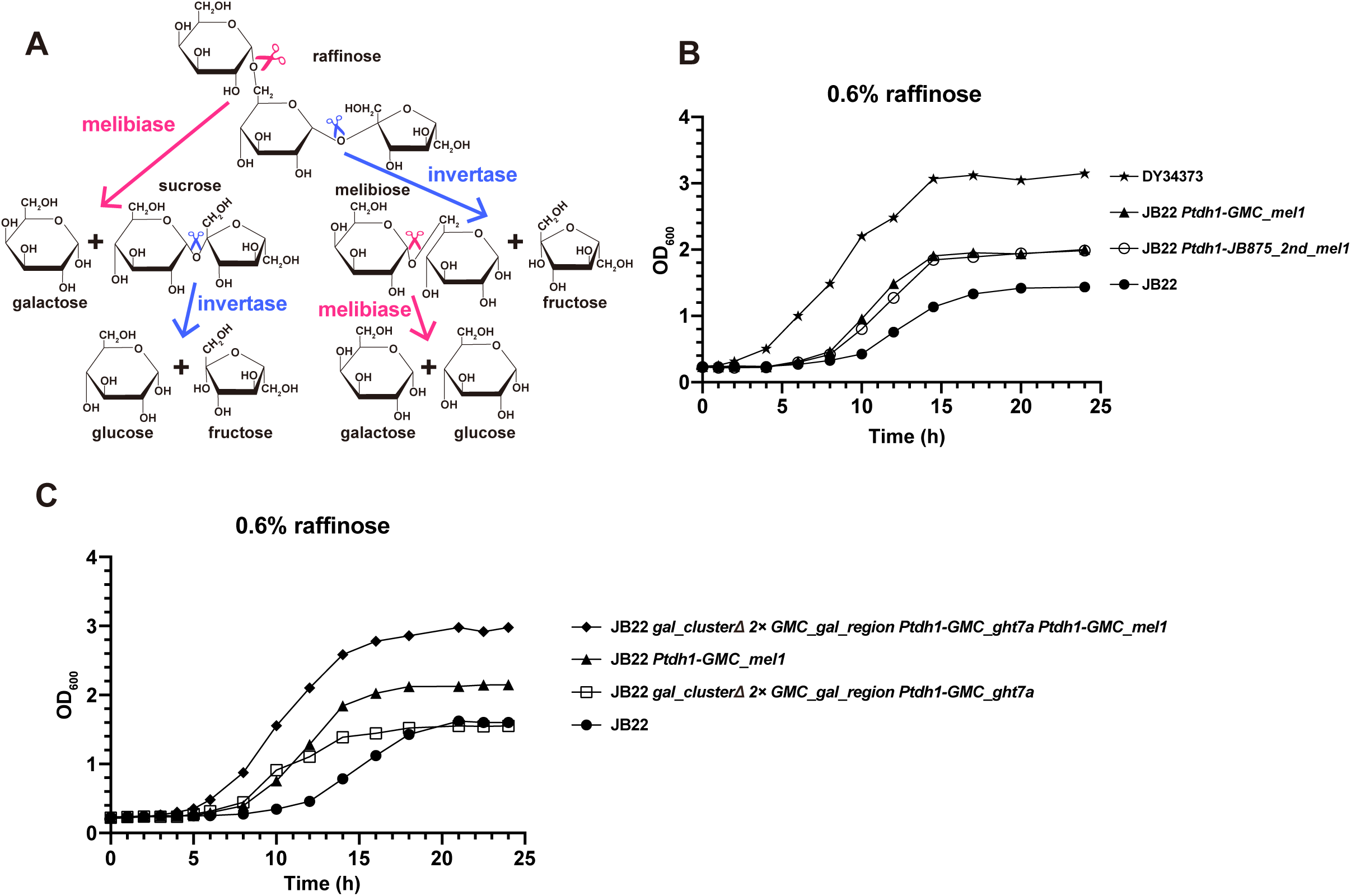
Strains that efficiently utilize melibiose and galactose can maximally utilize raffinose as a carbon source. (*A*) Schematic representation of the melibiase- and invertase-catalyzed hydrolysis reactions that convert raffinose into disaccharides and monosaccharides. (*B*) Growth curve analysis comparing the raffinose utilization abilities of JB22, JB22 expressing *GMC_mel1* under *Ptdh1*, JB22 expressing *JB875_2nd_mel1* under *Ptdh1*, and DY34373. (*C*) Growth curve analysis comparing the raffinose utilization abilities of JB22 and three JB22-derived strains.

We hypothesized that the Mel^+^ and Gal^F^ traits may confer a growth advantage in raffinose-rich environments and tested this idea (Fig. 7 *B* and *C*). We compared the growth of the Mel^−^ reference strain JB22 with two JB22-derived Mel^+^ strains—expressing either *GMC_mel1* or *JB875_2nd_mel1* under the *Ptdh1* promoter—and the naturally Mel^+^ and Gal^F^ strain DY34373 in a YE-based liquid medium containing raffinose as the sole carbon source (Fig. 7*B*). Under these conditions, biomass accumulation reflects the extent to which the constituent monosaccharides of raffinose are metabolized.

The JB22 culture reached a plateau OD_600_ of ∼1.5 after 20 hours. In contrast, both Mel^+^ derivative strains reached a higher plateau of ∼2.0 within 14 hours, indicating that melibiase expression enhances raffinose utilization. Notably, naturally Mel^+^/Gal^F^ isolate DY34373 achieved the highest biomass yield, reaching an OD_600_ of ∼3.1 after 14 hours—more than double that of JB22 (Fig. 7*B*).

We reasoned that the superior performance of DY34373 arises from its ability to not only hydrolyze raffinose via *GMC_mel1*-encoded melibiase but also to efficiently metabolize the released galactose through the multi-copy *GMC_gal_regions* and the functional galactose transporter *GMC_ght7a*. To evaluate this, we compared JB22 with three JB22-derived strains: one expressing *GMC_mel1* to gain melibiose utilization, one carrying two copies of the *GMC_gal_region* along with *Ptdh1*-driven *GMC_ght7a* to enhance galactose utilization, and one combining both genetic modifications (Fig. 7*C*).

The strain with improved galactose utilization but lacking melibiase did not surpass JB22 in final biomass, likely because it can only access the fructose component of raffinose through invertase-mediated hydrolysis. Its shorter lag phase may result from increased fructose uptake mediated by *GMC_ght7a*. The strain expressing *GMC_mel1* achieved higher biomass, consistent with its ability to utilize not only fructose but also glucose derived from melibiose breakdown. The strain possessing both melibiose hydrolysis ability and efficient galactose catabolism reached the highest biomass level, presumably because it can fully exploit all three monosaccharide units—fructose, glucose, and galactose—for growth (Fig. 7*C*).

These results demonstrate that the capacity to efficiently utilize melibiose and galactose confers a substantial growth advantage in raffinose, suggesting that the Mel^+^ and Gal^F^ phenotypes are adaptive traits that enhance fitness in raffinose-rich ecological niches.

## Discussion

In this study, we reveal that natural variation in galactose and melibiose utilization in *S. pombe* arises through an unexpected diversity of mechanisms—gene loss, repression, amplification, and HGT—demonstrating how microbial metabolic repertoires dynamically contract and expand during evolution.

Our identification of ethanol as an inducer of galactose utilization resolves a long-standing puzzle: why *S. pombe* possesses galactose utilization genes yet cannot grow in galactose liquid media. Ethanol similarly enhances glycerol utilization, suggesting a broader role in regulating non-preferred carbon metabolism. In *Sac. cerevisiae*, ethanol acts as a quorum-sensing molecule inducing filamentous growth, though its sensing mechanism remains unclear (48–50). Future work should explore whether ethanol acts via conserved signaling pathways.

While Gal^−^ phenotypes are rare in *Sac. cerevisiae* (1,51), they occur in 29% (17/58) of *S. pombe* isolates. Of these, 40% harbor *gal* cluster deletions; the remainder retain intact clusters but exhibit insufficient expression. Crucially, the results of our revertant screens implicate the Win1-associated MAPK pathway—known to respond to nutrient stress (52)—and the transcriptional repressor Ssn6 in repressing *gal* genes and causing the Gal^−^ phenotype. This parallels observations in *Sac. cerevisiae*, where *ssn6* mutations enable utilization of mannitol and sorbitol as carbon sources (53,54). Transcriptional repression may be a widespread strategy for maintaining latent metabolic potential in fluctuating environments.

Our discovery of the *gal-mel* cluster (GMC)—a tandemly amplified, horizontally acquired gene cluster that enables both rapid galactose catabolism and melibiose utilization (Gal^F^ and Mel^+^ traits)—highlights how eukaryotic microbes can gain complex metabolic capabilities through the rare but impactful introduction of foreign metabolic modules. The independent emergence of a second functional melibiase gene (*JB875_2nd_mel1*) in a distinct lineage underscores the recurrent potential for metabolic expansion via HGT. Notably, both GMC and *JB875_2nd_mel1* reside in subtelomeric regions—hotspots for genomic plasticity and foreign DNA incorporation (55,56).

The elevated synonymous substitution rates we observe in the ∼120-kb region flanking the GMCs (Fig. S12) invite comparison with analogous observations in *Saccharomyces*, where *d_S_* elevation extending into *GAL*-flanking regions has been interpreted as evidence of long-term balancing selection maintaining divergent reference and alternative *GAL* versions within and across species (8,57–59). We note, however, an important architectural distinction: in *Saccharomyces*, reference and alternative *GAL* versions occupy the same genomic loci, whereas in *S. pombe* the GMCs reside near the right telomere of chromosome I while the canonical *gal* cluster is located on chromosome II, and all four GMC-containing isolates also retain the canonical *gal* cluster. This non-allelic relationship complicates a direct application of the balancing selection model to the GMCs. Nevertheless, balancing selection could still operate at the level of the chromosome I right end, maintaining two versions of this subtelomeric region—one harboring the GMCs and one without. The rarity of GMC-containing isolates (4 of 58 surveyed) is more consistent with the GMCs conferring a fitness advantage only in specific ecological niches.

The ecological relevance of the Gal^F^ and Mel^+^ traits is underscored by our finding that efficient galactose and melibiose utilization confers a substantial growth advantage on raffinose—a plant-derived trisaccharide abundant in certain natural niches. Notably, two of the four GMC-containing strains were isolated in 1957 from fermentation vessels at the Leningrad (now St. Petersburg) Hydrolysis Plant (60), where ethanol was produced by yeast fermentation of acid-hydrolyzed wood biomass. Wood hydrolysates contain a mixture of sugars, including hemicellulose-derived galactose, and thus could plausibly favor strains with enhanced galactose utilization capacity. A third strain was also isolated in the Soviet Union no later than 1958, but its exact origin is unclear. The fourth strain was isolated from grapes in southern Ukraine. Whether the grape isolate represents a feral strain that escaped from an industrial environment is an open question.

In summary, we have uncovered a rich landscape of metabolic variation in *S. pombe*, driven by a combination of reductive (gene loss, repression) and expansive (amplification, HGT) processes. Our findings highlight that microbial adaptation is not driven solely by selection on standing genetic variation—it is propelled by rare, impactful innovations that reshape metabolic potential.

## Materials and Methods

### Fission yeast strains and plasmids

*S. pombe* strains used in this study are detailed in *SI Appendix*, Dataset S1. Plasmids employed for strain construction are documented in *SI Appendix*, Dataset S2. Genome sequencing data have been deposited in the NCBI SRA and China National GeneBank (CNGB), and PacBio HiFi genome assemblies have been deposited in GenBank and CNGB (accession numbers are provided in *SI Appendix*, Dataset S1). The sequence and annotation of the GMC (*gal-mel* cluster) are archived in GenBank under accession number PV988365. The sequence and annotation of nine additional genes (*JB875_2nd_mel1*, *GMC_DUF999b*, *DY39827_DUF999-1*, *DY39827_DUF999-2*, *DY39827_DUF999-3*, *DY39827_DUF999-4*, *DY39827_DUF999-5*, *DY39827_DUF999-6*, and *DUF999-11*) have been deposited in GenBank under accession numbers PV988366-PV988374.

Fission yeast methods, including transformation, crosses, tetrad dissection, and iodine staining analysis of sporulation, were performed as described by Forsburg and Rhind (2006) (61). Gene deletions were primarily achieved through homologous recombination. Typically, about 300 bp sequences upstream and downstream of the target gene’s coding region were used as homology arms. For genes that were difficult to delete, homology arms of 500 bp to 1 kb were employed. The coding region was replaced with an antibiotic resistance marker or an auxotrophy-complementing marker.

To convert homothallic strains into heterothallic strains and mark the mating type locus, the *mat1Δ17* mutation was constructed by replacing the 140 bp sequence between the H1 homology box and the nearby SspI restriction site at the *mat1* locus with the *natMX* resistance marker (62,63). Nourseothricin-resistant colonies were replica-plated onto SPA plates and stained with iodine. Colonies that failed to stain with iodine were streaked onto YES solid media. After two rounds of streaking, iodine staining and microscopic observation were used to verify the absence of spore production.

We employed the HO endonuclease to delete the GMC array in DY39827. This method is similar to the inducible telomere formation system reported by Wang et al. (64), except that HO is used instead of I-SceI to generate a double-strand break (DSB). A DNA sequence containing telomeric repeats, an HO cleavage site, and the *natMX* resistance marker was integrated at the centromere side of the GMC array. Upon HO expression, cells that repair the DSB through telomere formation are expected to lose the *natMX* resistance marker and the GMC array. Using this approach, we successfully obtained a DY39827 strain lacking GMCs, although the deleted region was slightly larger than anticipated.

The *gal7Δ::mECitrine* reporter strains were constructed by replacing the *gal7* coding sequence with the coding sequence of mECitrine and the *kanMX* marker, as previously described (65–67).

All plasmids used in this study were constructed by Gibson Assembly, combining purified PCR products with linearized vectors generated by restriction digestion.

### Spot assay for phenotypic analysis

Strains were activated on YES solid media, cultured overnight in YES liquid media at 30 °C. After dilution to logarithmic phase, cells were washed and 0.3 OD_600_ units of cells were resuspended in 1 ml of water. Serial 5-fold dilutions were performed in a 96-well plate, starting with 200 μl of cell suspension and transferring 40 μl between adjacent wells.

Before spotting, a 48-pin frogger was sterilized by immersion in 95% ethanol and flamed three to four times. After cooling, the frogger was dipped into the 96-well plate. The plate was held steady while the frogger was gently agitated to ensure even coating of the pins. The frogger was then lifted vertically and carefully pressed onto the solid agar surface. After approximately 10 seconds, the frogger was lifted vertically again. The procedure was repeated if contamination or incomplete spotting occurred.

Culture plates were inverted and incubated at 30 °C. On YES media, strains typically grew well within 2 to 3 days. On media containing galactose (TCI, G0008) or glycerol (Beijing Yili Fine Chemicals Co., Ltd.), growth was visible after about 4 days, with obvious growth achieved by 6 to 7 days. On melibiose-containing media, strains capable of utilizing melibiose (TCI, M0050) generally grew well within 3 to 4 days.

To prevent ethanol volatilization, culture plates supplemented with ethanol were wrapped in parafilm. Notably, for solid culture media containing melibiose, if a strain can utilize melibiose, the melibiose in the media will be hydrolyzed by the extracellular melibiase secreted by the strain, producing monosaccharides that diffuse throughout the solid culture media. On the same media, strains that cannot utilize melibiose may utilize the monosaccharides diffused from the surrounding culture media, leading to a misleading appearance of growth. To prevent this phenomenon, gaps were created on the agar plates by cutting out agar slices after spotting.

### Growth curve assays in liquid media

For growth in galactose or glycerol (with or without 0.5% ethanol), strains were pre-cultured in 2% glycerol with 0.5% ethanol media and cultured overnight. Harvested 0.1 OD_600_ units during logarithmic phase, washed, and transferred to 20 ml of media containing galactose or glycerol (with or without ethanol). Growth was monitored by measuring OD_600_ at different time points to construct growth curves.

To assess growth in raffinose (Sigma, R0250), strains were grown in YES liquid media to logarithmic phase. 10 OD_600_ units of cells were collected, washed, and transferred to 50 ml of media containing raffinose. OD_600_ values were measured at various time points to generate growth curves.

### Quantitative PCR (qPCR) for copy number analysis

Primers were designed using NCBI’s Primer-BLAST tool. Genomic DNA was extracted and quantified using a Qubit fluorometer.

qPCR was performed in 20-μl reactions containing 10 μl of 2× SYBR qPCR Master Mix (without ROX), 2 μl of template, 0.4 μl of each primer, and 7.2 μl of deionized water. The reaction program included an initial denaturation step at 95 °C for 3 min, followed by 45 cycles of 95 °C for 10 s and 55 °C for 50 s. Melting curve analysis was performed using the “Quick plate 96 wells-SYBR only.pltd” setting.

A standard curve was generated using serial dilutions of the DNA from a control strain containing a single copy of *GMC_gal1* integrated into the genome. The copy number of *GMC_gal1* in each sample was calculated using the standard curve.

### Microscopy observation and fluorescence quantification

Live cell imaging was performed using a DeltaVision PersonalDV system equipped with mCherry, YFP, and CFP filters and a 100× 1.4 NA objective. Images were captured using a Photometrics CoolSNAP HQ2 camera and analyzed using SoftWoRx software. For fluorescence quantification, cell boundaries were segmented using Cellpose (version 2.1.1) (68), and the mean fluorescence intensity for each sample was calculated using the LabelsToRois plugin in Fiji (69,70).

### Tn5 transposase-based library preparation for Illumina sequencing

Genomic DNA was extracted from each sample using the MasterPure Yeast DNA Purification Kit. Prior to phenol-chloroform extraction, 1 μl of RNase was added, and the samples were incubated at 37 °C overnight for RNA digestion.

Tn5 transposase and matched adapters were assembled in a 20-μl reaction containing 8 μl of MEDS-A+B (adapters), 6 μl of Tn5 transposase, and 6 μl of deionized water. The mixture was incubated at 25 °C for 1 hour. Then, tagmentation reactions were performed in a 20-μl reaction containing 2 μl of the assembled Tn5, 1 μl of genomic DNA, 4 μl of 5× TAPS-DMF reagent, and 13 μl of deionized water. The reaction was carried out at 55 °C for 1 hour. The labeled products were purified using the Illustra GFX PCR DNA and Gel Band Purification Kit to inactivate Tn5 transposase.

Indexing PCR was performed to enrich the tagmented DNA. The 50-μl reaction mixture consisted of 25 μl of 2× buffer from the KAPA HiFi HotStart ReadyMix reagent, 1 μl of each custom-designed index primer, 4 μl of the purified tagmentation reaction product, and 19 μl of deionized water. The PCR program included an initial step at 72 °C for 3 minutes and 95 °C for 30 seconds, followed by 14 cycles of 95 °C for 10 seconds, 55 °C for 30 seconds, and 72 °C for 30 seconds. The PCR products were verified by electrophoresis. If the expected product size was correct, target fragments were isolated using solid-phase reversible immobilization (SPRI) beads and sent to a commercial sequencing company for next-generation sequencing. Paired-end sequencing (150 bp) was performed using the Illumina NovaSeq 6000 platform, generating 2 Gb of raw data per sample.

### Spontaneous Gal^+^ revertant screens

5 Gal^−^ strains with an intact *gal* cluster (JB878, JB762, JB848, DY39825, and JB852), 1 Gal^−^ strain harboring a naturally occurring *gal* cluster deletion (JB874), and 1 reference strain with the *gal* cluster experimentally deleted were cultured in YES liquid media overnight at 30 °C. Diluted once, 1 OD_600_ unit of cells was collected during logarithmic phase, serially diluted and plated on YES solid media. Plates were incubated at 30 °C for three days to obtain numerous single colonies.

For each strain, ten replicates were prepared. In each of ten 1.5 ml microcentrifuge tubes, 100 μl of deionized water was added. An appropriate number of single colonies (4-5 for larger colonies or 10 for smaller colonies to ensure that the amount of cells exceeds 1 OD_600_ unit) were picked and resuspended in the water. The cell suspensions were washed three times with water and resuspended in 1 ml of water. Each suspension was diluted 5×, and the concentration was measured. Finally, 1 OD_600_ unit of cells from each sample was spread onto YE + 2% galactose + 0.5% ethanol solid media, with each plate representing one pool. To prevent ethanol volatilization, plates were wrapped with parafilm.

Plates were incubated at 30 °C for several days (fast-growing strains formed small colonies within four days, while slow-growing strains required over two weeks). A single large colony was selected from each plate and streaked onto fresh YES solid media. Illumina sequencing libraries were prepared using the Tn5 library construction method. The resulting libraries were sent for Illumina sequencing.

Raw reads from the Illumina NovaSeq 6000 platform (PE150) were quality-filtered using fastp (v0.20.0, https://github.com/OpenGene/fastp) with a minimum length cutoff of 70 bp. Clean reads of the Gal^+^ mutant clones were mapped to the reference genome using BWA (v0.7.18-r1188, https://github.com/lh3/bwa) (71). Subsequent processing employed Picard (v2.17.11, “Picard Toolkit.” 2019. GitHub Repository. https://broadinstitute.github.io/picard/) tools for coordinate sorting (SortSam) and duplicate marking (MarkDuplicates). For variant detection, we performed parallel calling with bcftools (v1.9, https://github.com/samtools/bcftools) (72), and DeepVariant (v1.3.0, https://github.com/google/deepvariant) (73) to identify single nucleotide polymorphisms (SNPs), insertions and deletions (INDELs), and multiple nucleotide polymorphisms (MNPs) (74). Resulting variants were decomposed into allelic primitives using vcfallelicprimitives (MNP splitting) and annotated with snpEff (v4_3t, https://pcingola.github.io/SnpEff/). Information about sequencing data at positions of all variants was obtained using bam-readcount (https://github.com/genome/bam-readcount) with the following parameters: --min-mapping-quality 20 --min-base-quality 20 (75). Based on the bam-readcount results, the reference allele frequency was calculated. Using snpEff-generated variant impact annotation and reference allele frequency thresholds (SNP < 0.15, INDEL < 0.25), we selected a total of 355 non-silent variants.

The following filtering criteria were further applied to the 355 variants: (1) repeat region exclusion using GenMap (v1.3.0, https://github.com/cpockrandt/genmap) with the following parameters: -k 100 – E 2 (76), (2) exclusion of dubious genes and *S. pombe*-specific genes, (3) background variant filtering (≥ 4 samples), (4) depth filtering (≥ 10), (5) exclusion of mitochondrial variant sites, (6) manual inspection of the variants. This yielded 34 high-confidence variants (listed in *SI Appendix*, Dataset S3). The analysis scripts are available in a GitHub repository (https://github.com/DuXiaomin-NIBS/research-scripts.git).

Illumina read mapping data were also utilized for copy number variant (CNV) analysis. Mean read depths were calculated for: (1) all essential genes on chromosome I, and (2) 1-kb genomic windows across all Gal^+^ mutant clones using SAMtools (v1.10, https://github.com/samtools/samtools) (77). Gene copy numbers were determined through depth-ratio normalization.

### SNP pattern analysis of the chromosome II *gal* cluster region in 58 *S. pombe* natural isolates

Raw sequencing reads generated from the Illumina NovaSeq 6000 System (PE150) were quality filtered using fastp. To further enhance data quality, KAT (v2.4.2, https://github.com/TGAC/KAT) was used to eliminate read pairs containing low-frequency k-mers (78). The resulting high-quality reads from all 58 *S. pombe* natural isolates were mapped to a modified reference genome that included the GMC sequence using BWA (v0.7.18-r1188, https://github.com/lh3/bwa) (71). Subsequent processing followed the pipeline described in the previous section, utilizing Picard tools for BAM file sorting and duplicate read marking. HaplotypeCaller, a tool in GATK (v3.8-1-0-gf15c1c3ef, https://github.com/broadinstitute/gatk), was used with the following parameters: --emitRefConfidence GVCF --variant_index_type LINEAR --variant_index_parameter 128000 -mbq 20 -stand_call_conf 20 (79,80). Individual variant call format (VCF) files from all 58 natural isolates were merged for joint variant calling. We filtered identified SNPs using GATK’s SelectVariants tool with stringent criteria: --filterExpression “QD < 2.0 || FS > 60.0 || MQ < 40.0” --filterName “my_snp_filter” -G_filter “DP<10” -G_filterName “lowiDP”. Genotypes failing these filters were designated as no-call and excluded from subsequent analysis. The identical pipeline was applied for INDEL variant detection.

The analysis focused on the chromosome II region spanning positions 4467518 to 4492116. Following the extraction and merging of quality-passed SNP and INDEL sites, multi-allelic loci were removed prior to visualization using a customized R script that generated comprehensive heatmap representations. The analysis scripts are available in a GitHub repository (https://github.com/DuXiaomin-NIBS/research-scripts.git).

### Long-read genome sequencing

Strains were revived on YES solid media, cultured in YES liquid media overnight at 30 °C, and expanded to 1000 ml of YES medium. Cells were harvested at OD_600_ = 0.8, 1000 OD_600_ units were collected and immediately flash-frozen using liquid nitrogen.

The frozen cell pellets were subsequently transferred to Wuhan Frasergen Bioinformatics Co., Ltd for high-molecular-weight genomic DNA isolation and sequencing library preparation. Two types of long-read sequencing platforms were employed: (1) Pacific Biosciences (PacBio) using the Sequel platform and (2) high-fidelity PacBio HiFi using the Sequel II platform.

### Genome assembly and phylogenetic analysis

We acquired short-read sequencing data for all 58 *S. pombe* natural isolates and seven fission yeast species (*S. pombe*, *S. octosporus*, *S. lindneri*, *S. osmophilus*, *S. cryophilus*, *S. japonicus*, and *S. versatilis*) through our own sequencing efforts and published datasets (18,21,81–84). Within the natural isolate collection, 18 strains possess PacBio long-read sequencing data (83), while 3 strains (DY39827, JB852, and JB875) have high-fidelity PacBio HiFi data. All seven fission yeast species also have corresponding PacBio HiFi sequencing data available.

For genome assembly, we implemented the following pipelines: (1) Short-read sequencing data: A5-miseq (v20160825) with default parameters (85). (2) PacBio long-read sequencing data: Parallel assembly using Canu (v1.8, https://github.com/marbl/canu) (86), Flye (v2.6, https://github.com/mikolmogorov/Flye) (87), and Raven (v1.8, https://github.com/lbcb-sci/raven) (88). (3) PacBio HiFi sequencing data, Parallel assembly using canu (v2.2, https://github.com/marbl/canu) (89), and hifiasm (v0.16.1, https://github.com/chhylp123/hifiasm) (90).

All generated and published genome assemblies served as the subjects in BLAST. Sequences were extracted via tBLASTn, spanning from initiation to termination codons, followed by translation to amino acid sequences. We removed redundant sequences using CD-HIT (v4.6.2, https://github.com/weizhongli/cdhit/releases) with the following parameters: -c 1 -n 5 -G 1 -g 1 -b 20 -s 0.0 -aL 0.0 -aS 0.0 -S 0 -d 0 (91). These sequences were combined with orthologous sequences identified through BLASTp searches against non-fission yeast species. Multiple sequence alignment was conducted using MUSCLE (v3.8.31, https://github.com/rcedgar/muscle) with default settings (92). For phylogenetic tree construction, we employed IQ-TREE2 (v2.1.4, https://github.com/iqtree/iqtree2) under a maximum likelihood (ML) framework (93). The software was set to automatically select the optimal substitution model. Branch support assessment was performed using 1,000 bootstrap replicates and 1,000 SH-aLRT tests. Each analysis was repeated 10 times to identify the optimal tree topology. FigTree (v1.4.3, https://github.com/rambaut/figtree/releases) was used to visualize the final phylogenetic trees.

The neighbor-joining (NJ) tree based on SNPs was constructed using MEGA (v11.0.13) with the Jukes-Cantor substitution model (94). SNP and INDEL variants were identified following the methodology detailed in a previous section. Topological robustness was evaluated through bootstrap resampling with 1,000 replicates.

The evolutionary relationships among the seven fission yeast species were determined using complete and single-copy BUSCO genes. Ortholog identification was performed using BUSCO (v5.4.7; https://busco.ezlab.org/) with the ascomycota_odb10 dataset, revealing 1,128 complete and single-copy BUSCO genes present across all seven species (95). The phylogenetic tree construction pipeline comprised five sequential steps: (1) Extraction of all complete and single-copy BUSCO genes shared among the seven fission yeast species. (2) Multiple sequence alignment of protein sequences using MAFFT (v7.471, https://github.com/GSLBiotech/mafft) with default parameters (96). (3) Resulting alignments were trimmed with trimAL (v1.4.rev15, https://trimal.cgenomics.org) to remove low-quality alignment regions (97). (4) Processed alignments were concatenated into a supermatrix using AMAS (v1.0, https://github.com/marekborowiec/AMAS/) (98). (5) A maximum likelihood (ML) tree was generated using IQ-TREE2.

### Copy number analysis of *JB875_2nd_mel1*

Paired-end sequencing reads (150 bp) of JB875 from the Illumina NovaSeq 6000 platform were quality-filtered using fastp. Processed reads were mapped to the JB875 PacBio HiFi genome assembly using BWA (v0.7.18-r1188, https://github.com/lh3/bwa) (71). Mean read depths were calculated for: (1) all essential genes on chromosome I, and (2) 5-kb genomic windows across all JB875 chromosomes using SAMtools (v1.10, https://github.com/samtools/samtools) (77). Gene copy numbers were determined through depth-ratio normalization, comparing 5-kb window depths against essential gene depths using a customized R script, which simultaneously generated read depth plots. The analysis scripts are available in a GitHub repository (https://github.com/DuXiaomin-NIBS/research-scripts.git).

### Sliding-window analysis of nucleotide diversity

To compare nucleotide diversity between the reference genome and four GMC-containing strains, we performed sliding-window analysis using Simplot++ (v1.3, https://github.com/Stephane-S/Simplot_PlusPlus) with the following parameters: Step = 50, Window length = 1000 (99). The analytical workflow comprised three stages: (1) the *rnp24* to *gdt1* genomic region from all five strains (reference plus GMC-containing) was aligned using MAFFT (v7.453, https://github.com/GSLBiotech/mafft). (2) The resulting multiple sequence alignment was used as input for Simplot++. Pairwise nucleotide diversity plots were generated for all strain comparisons. (3) Annotation tracks derived from the alignment results were added to the diversity plots.

### Pairwise synonymous substitution rate analysis

To analyze sequence divergence, we calculated pairwise synonymous substitution rates using the following computational pipeline: (1) Coding sequences (CDS) and corresponding amino acid sequences were extracted for each gene pair. (2) An amino acid sequence alignment was generated using MAFFT (v7.453, https://github.com/GSLBiotech/mafft) with the G-INS-i algorithm (ginsi command). Using the resulting alignment as a guide, a codon-level nucleotide alignment was generated using PAL2NAL (v14, https://github.com/liaochenlanruo/PAL2NAL). (3) Biopython (v1.84) was used to compute non-synonymous (*d_N_*) and synonymous (*d_S_*) substitution rates via the codonseq.cal_dn_ds function (8,100,101).

### Synteny analysis and genomic region visualization

Syntenic relationships were visualized using the clinker web server (https://cagecat.bioinformatics.nl/tools/clinker) (102).

Gene positions were visualized at chromosomal resolution by inputting genomic coordinates into the chromoMap R package (v4.1.1, https://github.com/cran/chromoMap/blob/master/R/chromoMap.R), which generates chromosome-scale positional diagrams.

## Acknowledgments

We thank Guo-Song Jia for PacBio HiFi sequence assembly. We thank Jürg Bähler for making *S. pombe* natural isolates (JB strains) available through YGRC/NBRP, and YGRC/NBRP for providing strains. We are grateful to Dr. Vasily Bayraktar for providing information about the isolate he deposited at the NRRL Culture Collection (NRRL Y-48646). This work was supported by the National Key R&D Program of China 2024YFA0917400.

## Contributions

Conceptualization: X.-M.D. and L.-L.D.; Methodology and Investigation: X.-M.D., F.S. and L.-L.D.; Writing — Original Draft: X.-M.D. and L.-L.D.; Writing — Review and Editing: X.-M.D. and L.-L.D.; Funding Acquisition: L.-L.D.

## SI Appendix, Datasets

***SI Appendix*, Dataset S1 (separate file).** Strain list, including the *S. pombe* reference background strains and natural isolates used in this study, with information on the origins of homothallic natural isolates and accession numbers for both sequencing data and HiFi genome assemblies.

***SI Appendix*, Dataset S2 (separate file).** Plasmids used in this study for strain construction.

***SI Appendix*, Dataset S3 (separate file).** Gene duplications and mutations identified in spontaneous Gal^+^ revertant screens.

## Corresponding author

Correspondence to Li-Lin Du.

---

**Fig. S1.**
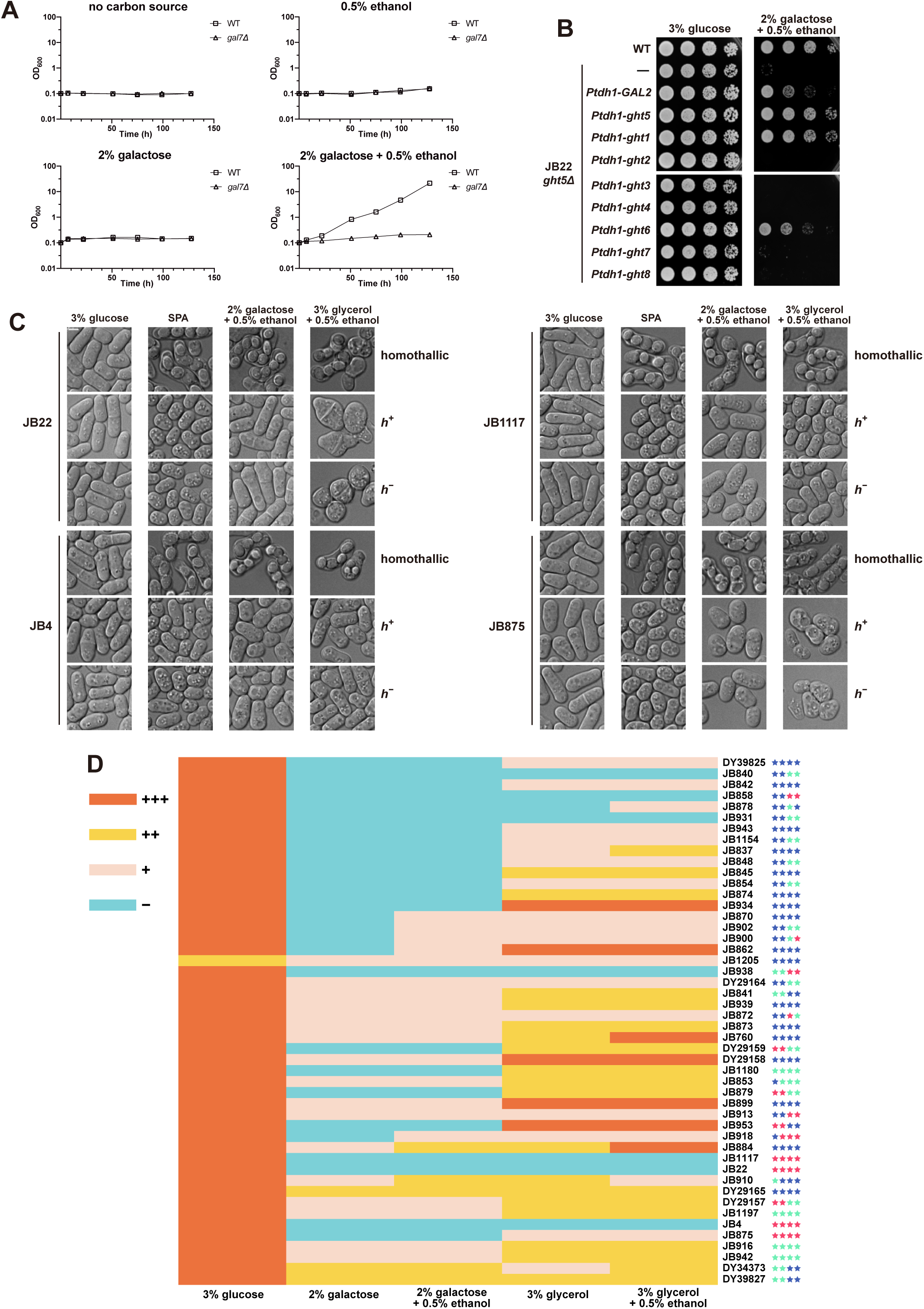
The roles of ethanol and *ght* genes on the galactose utilization ability of the *S. pombe* reference background strain, and the influence of self-mating ability on galactose utilization by *S. pombe* natural isolates. (*A*) The *S. pombe* reference background strain (JB22) cultured in liquid media can utilize galactose in the presence but not in the absence of ethanol. (*B*) Expressing Ght5 or Ght1 from the strong promoter *Ptdh1* fully rescued the galactose utilization defect of *ght5Δ*, while expressing Ght6 or the budding yeast galactose transporter Gal2 from the same promoter partially rescued the defect. (*C*) Microscopy observations of homothallic and heterothallic strains of the JB22, JB4, JB1117, and JB875 backgrounds on four different solid media. Homothallic but not heterothallic strains formed spores on the sporulation medium SPA and on YE-based galactose and glycerol media. Scale bar, 3 µm. (*D*) A heatmap showing the growth phenotypes of 48 distinct homothallic natural isolates of *S. pombe* on five different YE-based solid growth media. As in Fig. 1*H*, the growth phenotype was classified as no obvious growth (−), weak growth (+), moderate growth (++), or robust growth (+++). In addition, four columns of stars adjacent to strain names denote growth improvement upon conversion to heterothallic strains under four media conditions (left to right: 2% galactose, 2% galactose + 0.5% ethanol, 3% glycerol, and 3% glycerol + 0.5% ethanol). Blue stars indicate no growth improvement, green stars indicate moderate growth improvement, and pink stars indicate strong growth improvement.

**Fig. S2.**
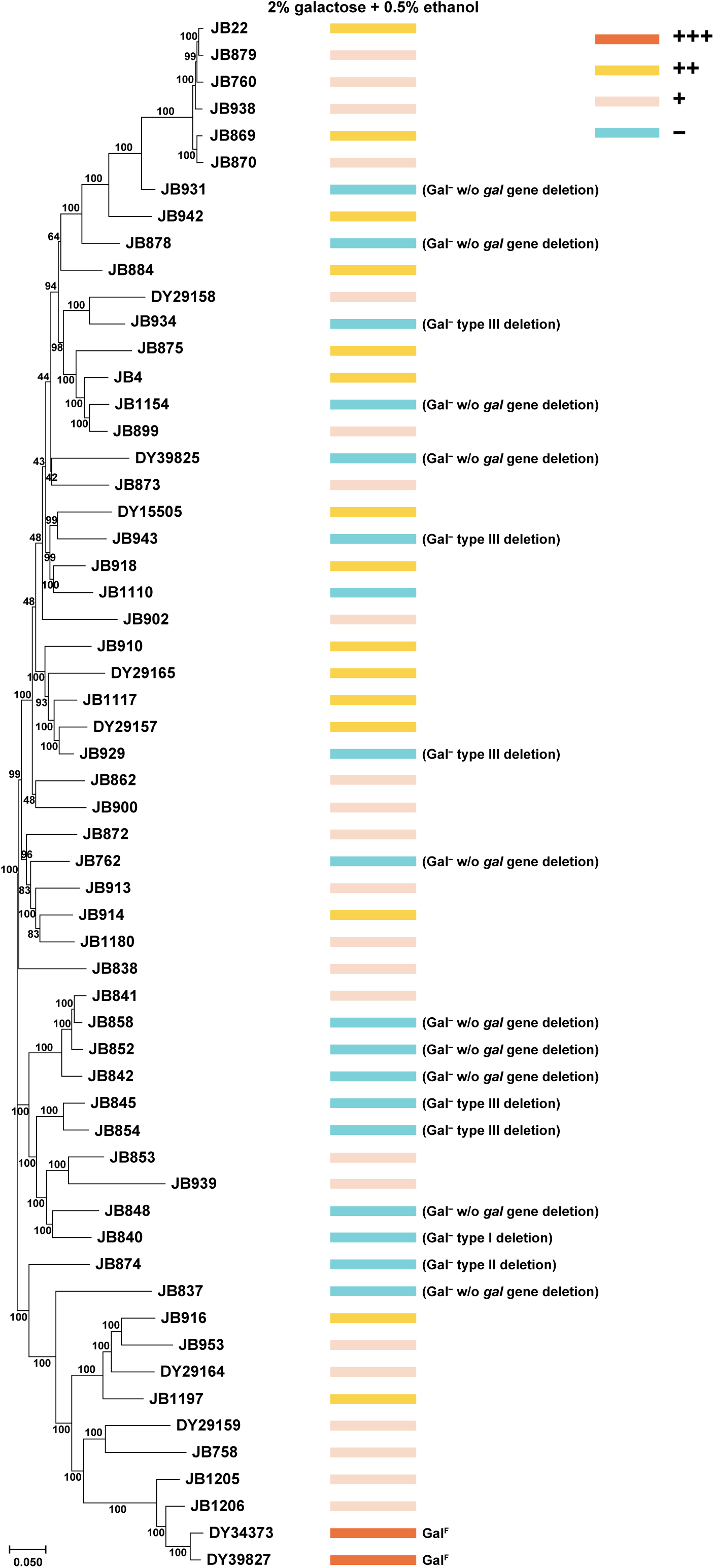
An SNP-based phylogenetic tree of the 58 *S. pombe* natural isolates used in this study. An SNP-based neighbor-joining (NJ) tree was constructed using MEGA and rooted by midpoint rooting. Branch support values were calculated by MEGA. The scale bar below tree indicates 0.05 substitutions per site. The growth phenotypes of naturally sterile or engineered heterothallic strains on 2% galactose + 0.5% ethanol are shown on the right.

**Fig. S3.**
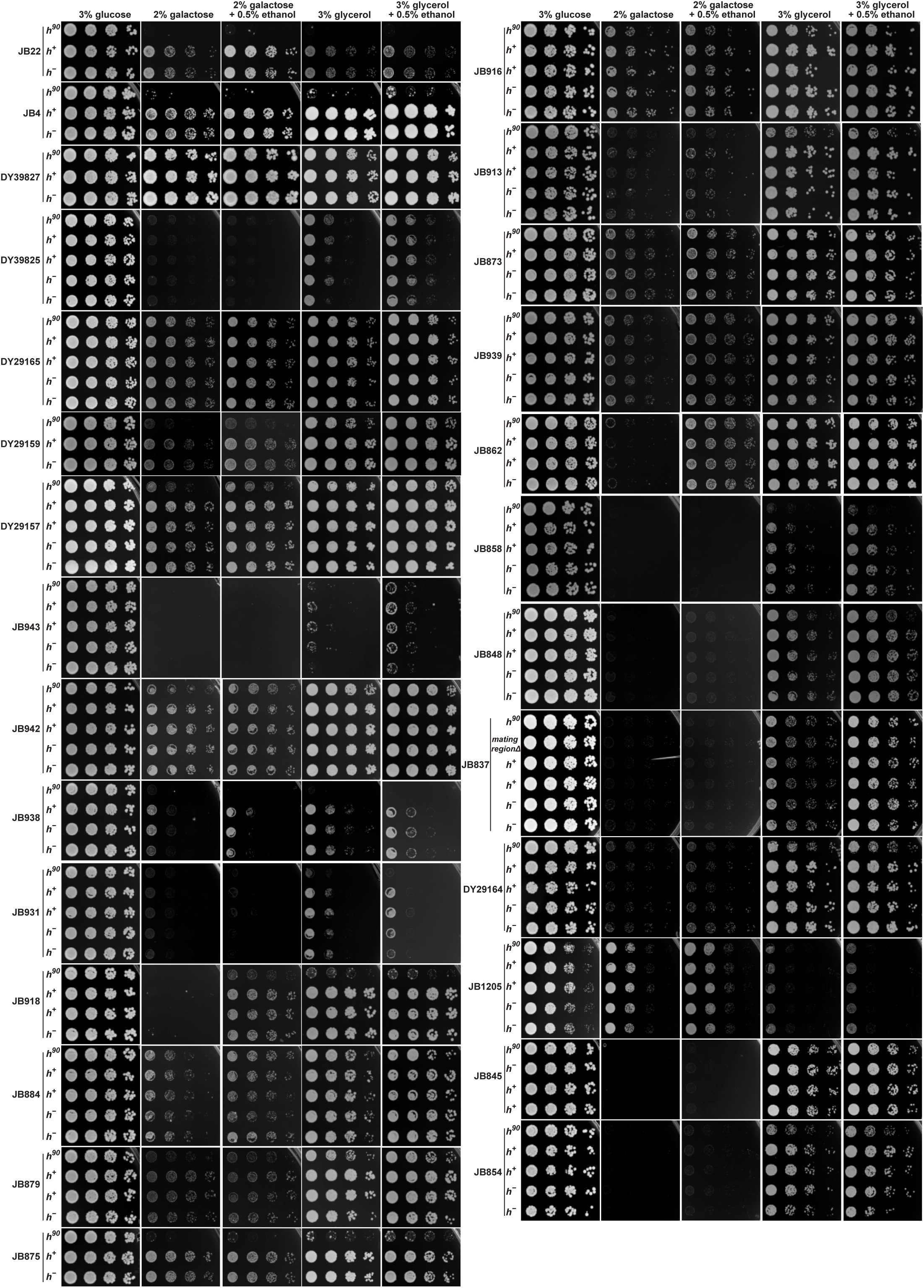
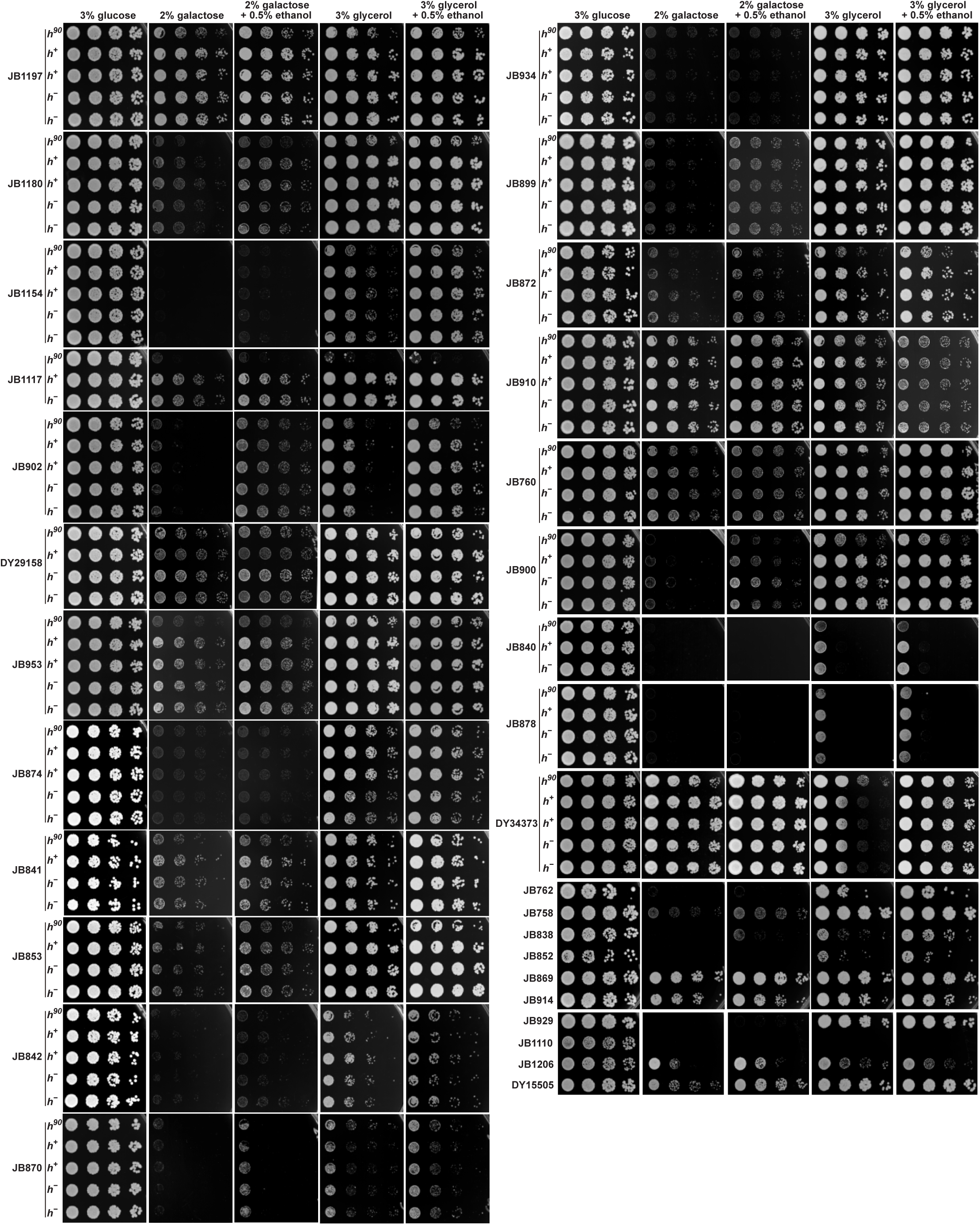
Growth phenotypes of 58 distinct *S. pombe* natural isolates on five types of YE-based solid media. The growth phenotype data shown here are the basis of the phenotype classifications shown in Fig. 1*H* and *SI Appendix*, Figs. S1*D* and S2. Images of the YE + 3% glucose plates were typically taken on day 3, while images of the other types of plates were typically taken on day 7. For brevity, homothallic strains are denoted as *h^90^*. The data for JB22 and JB4 were the same as those presented in Fig. 1*G*.

**Fig. S4.**
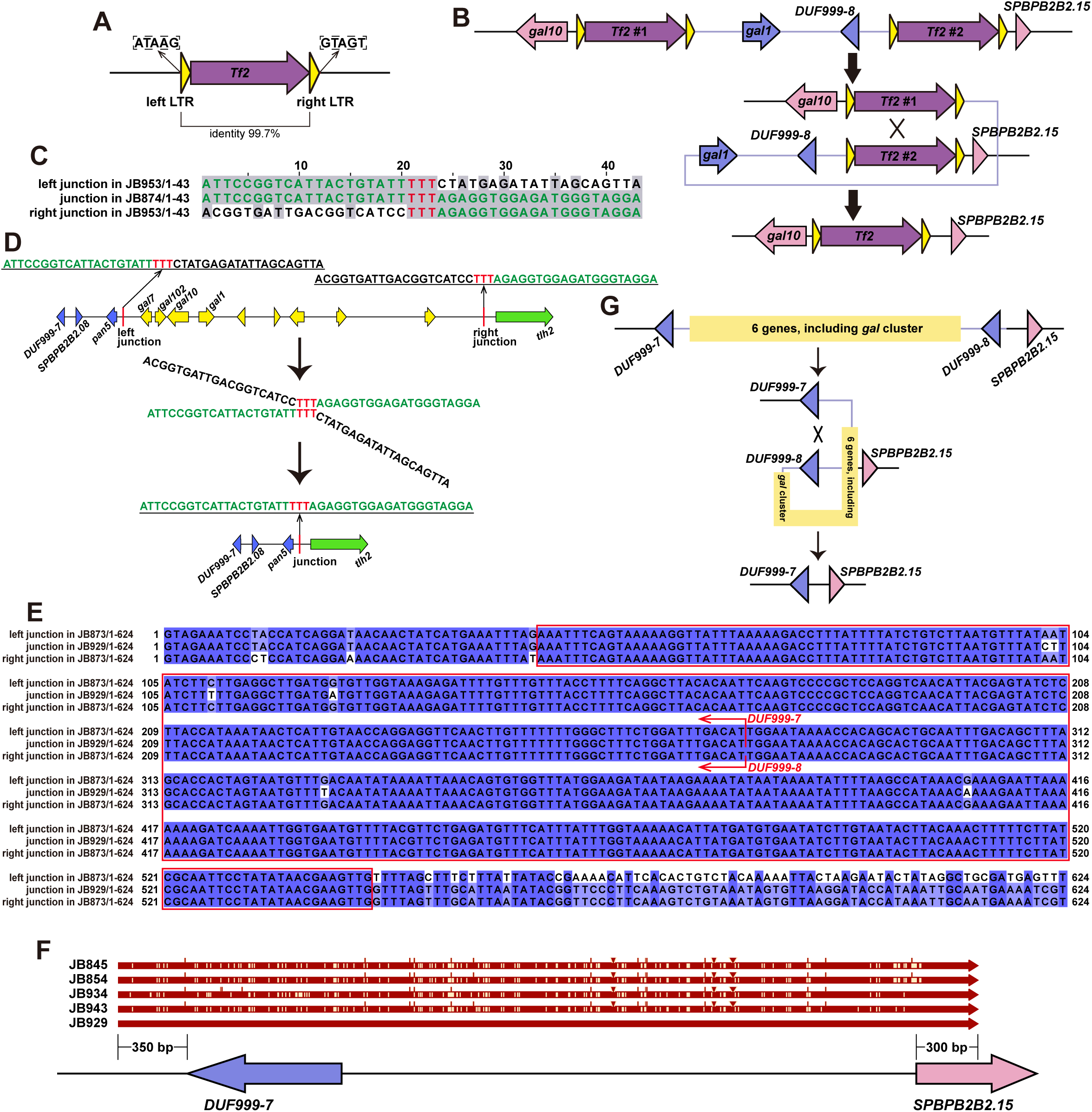
Details of three types of *gal* gene deletions. (*A*) Schematic diagram of the *Tf2* retrotransposon located in the *gal* cluster region of the type I deletion strain JB840. The sequence of JB840 was obtained from PacBio long-read genome assembly. The purple arrow represents the internal sequence of the *Tf2* retrotransposon, while the yellow triangles represent LTRs. The 5-bp sequences flanking the retrotransposon are shown. (*B*) A schematic depicting a possible scenario of how the deletion may have formed in the type I deletion strain. (*C*) A sequence alignment comparing the deletion junction in the type II deletion strain JB874 with the corresponding sequences at the left and right boundaries of the deleted region in a closely related strain lacking the deletion (JB953). The sequences of JB874 and JB953 were obtained from PacBio long-read genome assemblies. Identical bases are highlighted with a grey background. Sequences at the deletion breakpoint are highlighted in red font, while matched sequences on either side of the breakpoint are highlighted in green font. (*D*) A schematic depicting a possible process of how the deletion may have formed in the type II deletion strain. (*E*) A sequence alignment comparing the deletion junction in the type III deletion strain JB929 with the corresponding sequences at the left and right boundaries of the deleted region in a closely related strain lacking the deletion (JB873). Identical bases are highlighted with a blue background. The red box denotes the deletion breakpoint. The locations of the start codons of the two *DUF999* genes (*DUF999-7* and *DUF999-8*) are indicated with red bent arrows. (*F*) A diagram depicting the presence of an approximately 4.35-kb sequence spanning the deletion junction in the five type III deletion strains. The sequences of JB929, JB943, JB854, and JB873 were obtained from PacBio-based genome assemblies, while the sequences of the other two strains were from Sanger sequencing of PCR products. The diagram was generated using SnapGene. (*G*) A schematic depicting a possible scenario of how the deletion may have formed in type III deletion strains.

**Fig. S5.**
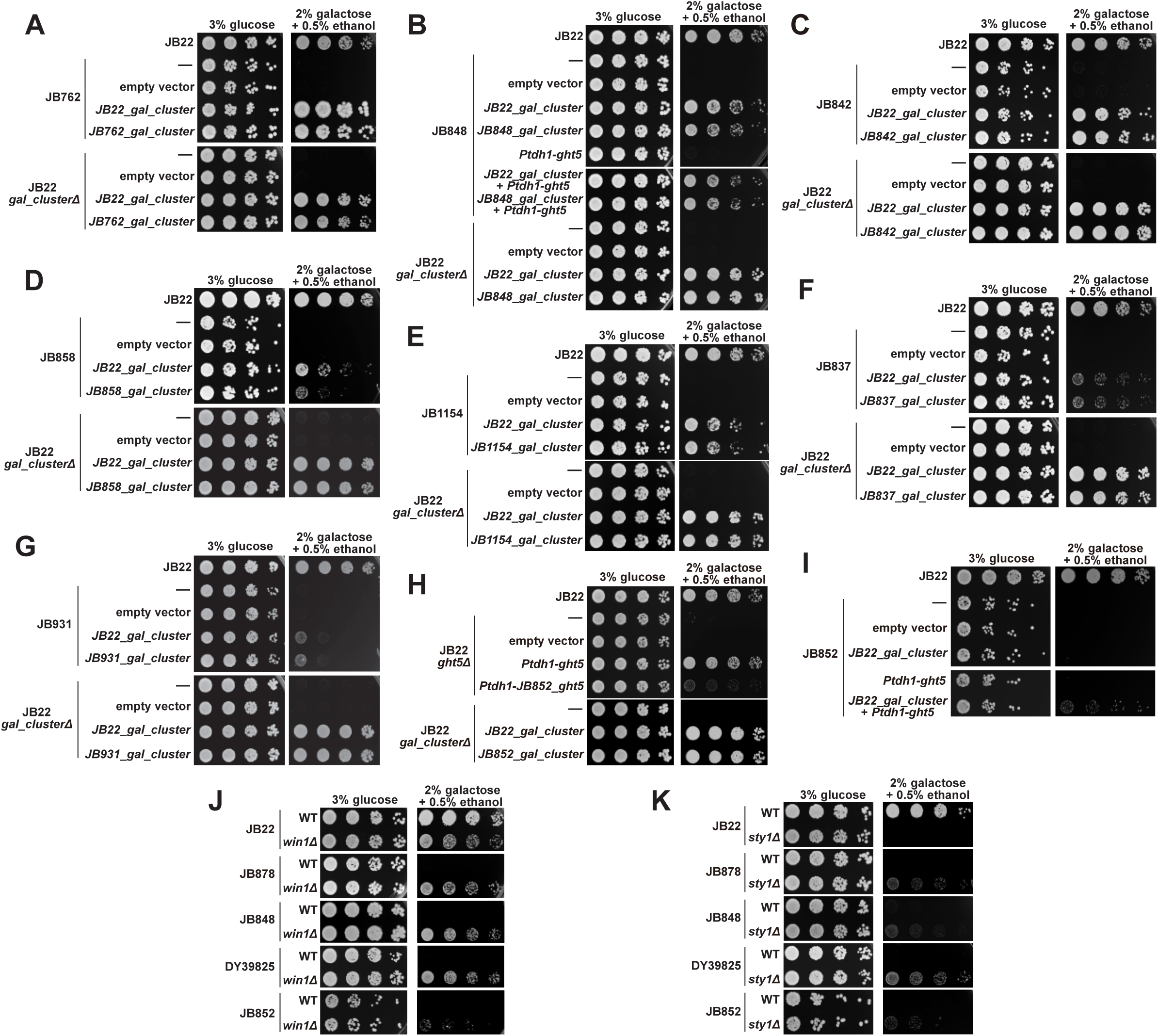
Characterization of Gal^−^ strains retaining an intact *gal* cluster. (*A-G*) The *gal*-gene-intact Gal^−^ strains JB762, JB848, JB842, JB858, JB1154, JB837, and JB931 can be converted from Gal^−^ to Gal^+^ through introducing one additional copy of the *gal* cluster. (*H*) The *gal* cluster, but not the *ght5* gene in JB852, is functional. The *ght5* alleles from JB852 and JB22 were expressed from the strong *Ptdh1* promoter. (*I*) JB852 was converted from Gal^−^ to Gal^+^ only when both the *gal* cluster and *ght5* from JB22 were introduced. (*J*) Deletion of *win1* weakened the galactose utilization ability of JB22 but converted JB878, JB848, DY39825, and JB852 from Gal^−^ to Gal^+^. The data for JB22 and JB878 were the same as those presented in Fig. 3*I*. (*K*) Deletion of *sty1* abolished the galactose utilization ability of JB22 but converted JB878, JB848, DY39825, and JB852 from Gal^−^ to Gal^+^.

**Fig. S6.**
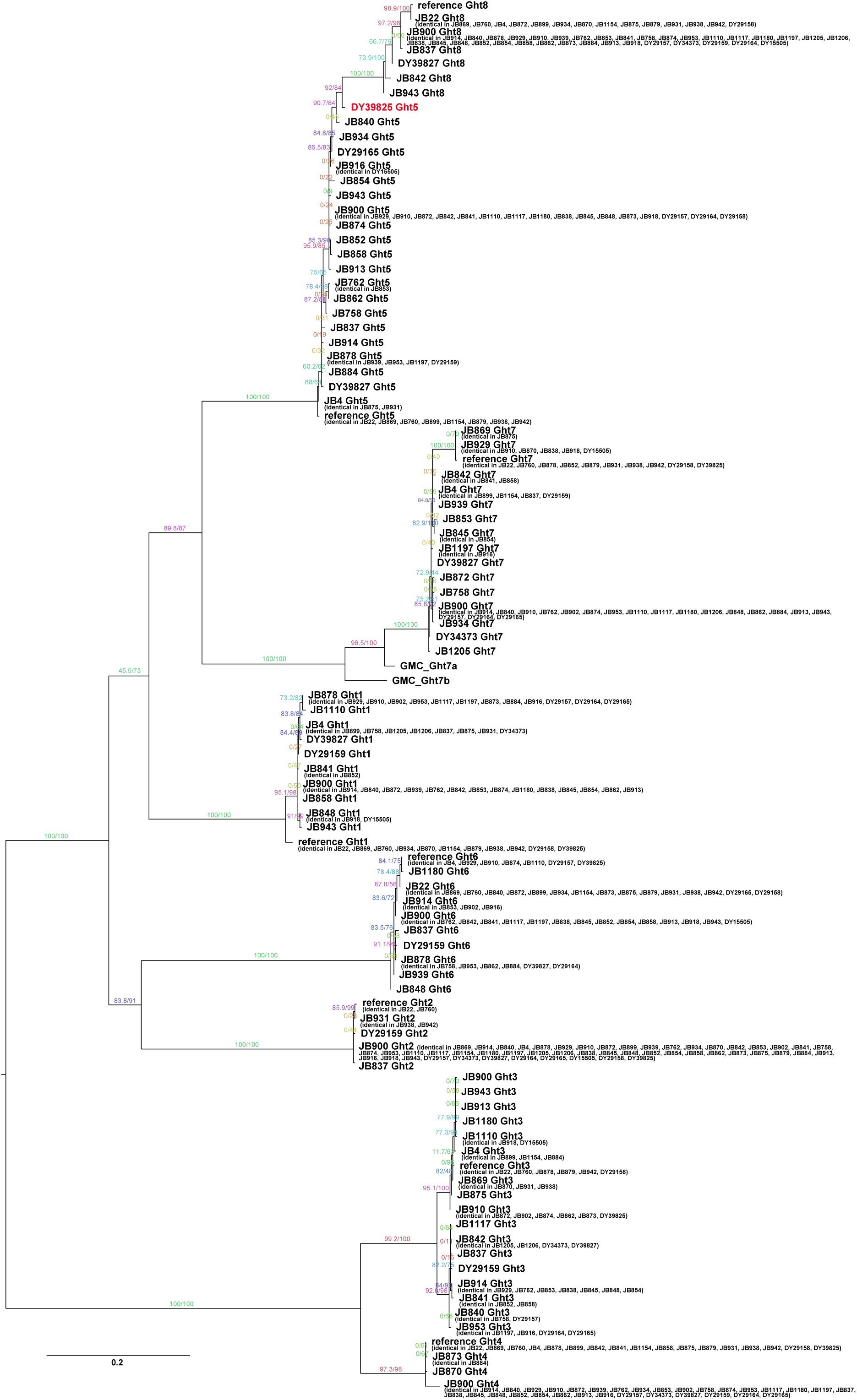
Phylogenetic analysis of hexose transporters (Ght family proteins) in 58 *S. pombe* natural isolates. A maximum likelihood (ML) tree of Ght family proteins from 58 natural isolates was constructed using IQ-TREE. The tree was rooted by midpoint rooting. Branch support values were calculated by IQ-TREE. The scale bar below tree indicates 0.2 substitutions per site. If a protein sequence is shared by multiple strains, only one sequence was used for tree construction and the corresponding node in the tree is labeled with the name of a representative strain, with other strains sharing the same sequence listed in parentheses. Ght5 of DY39825 is highlighted in red.

**Fig. S7.**
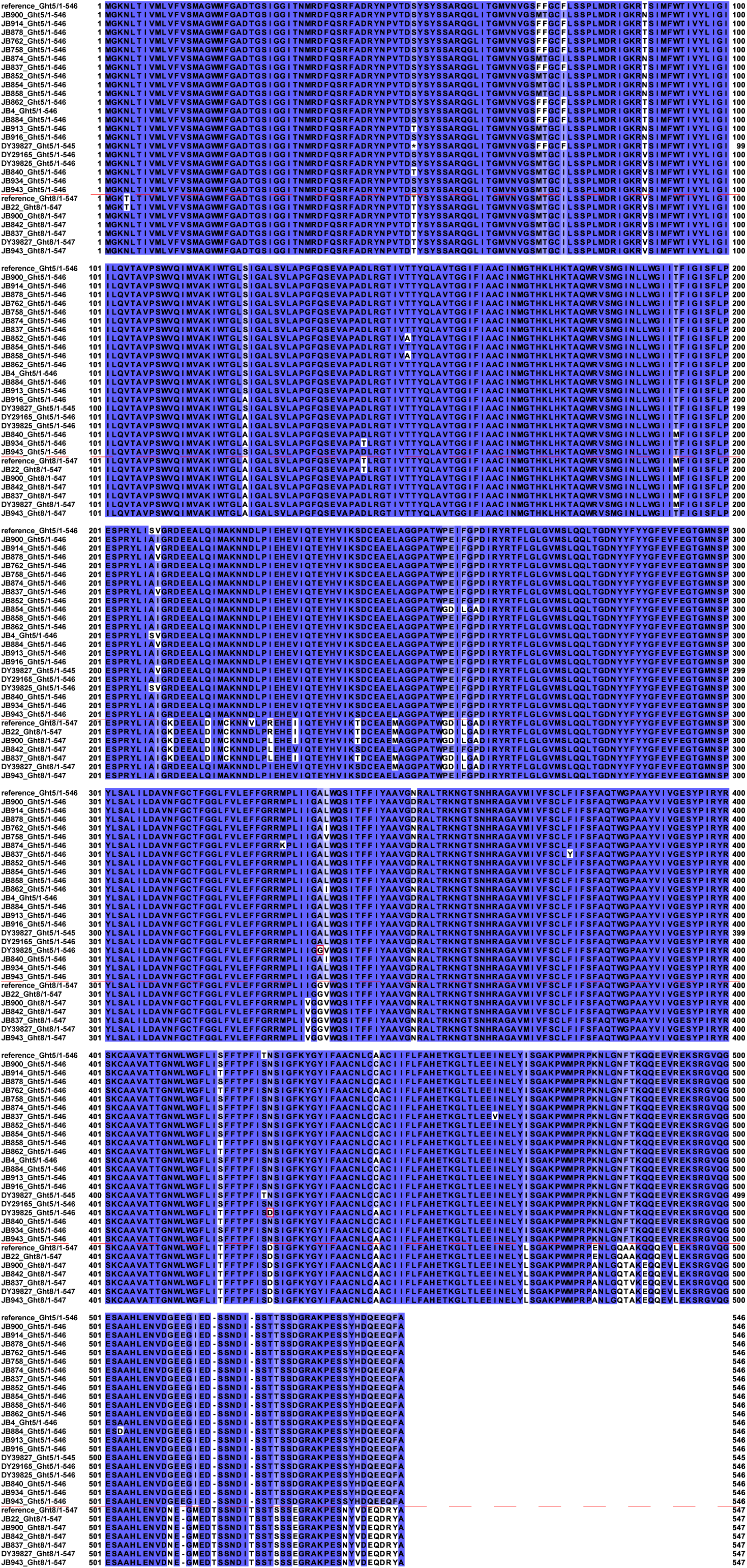
A sequence alignment comparing sequences of Ght5 and Ght8 in 58 *S. pombe* natural isolates. Identical bases are highlighted using blue background. Two red boxes denote the two amino acids indicative of recombination in the sequence of Ght5 of DY39825. A red dashed line separates Ght5 sequences from Ght8 sequences. For sequences shared by multiple strains, only the name of the representative strain shown in *SI Appendix*, Fig. S6 is given here.

**Fig. S8.**
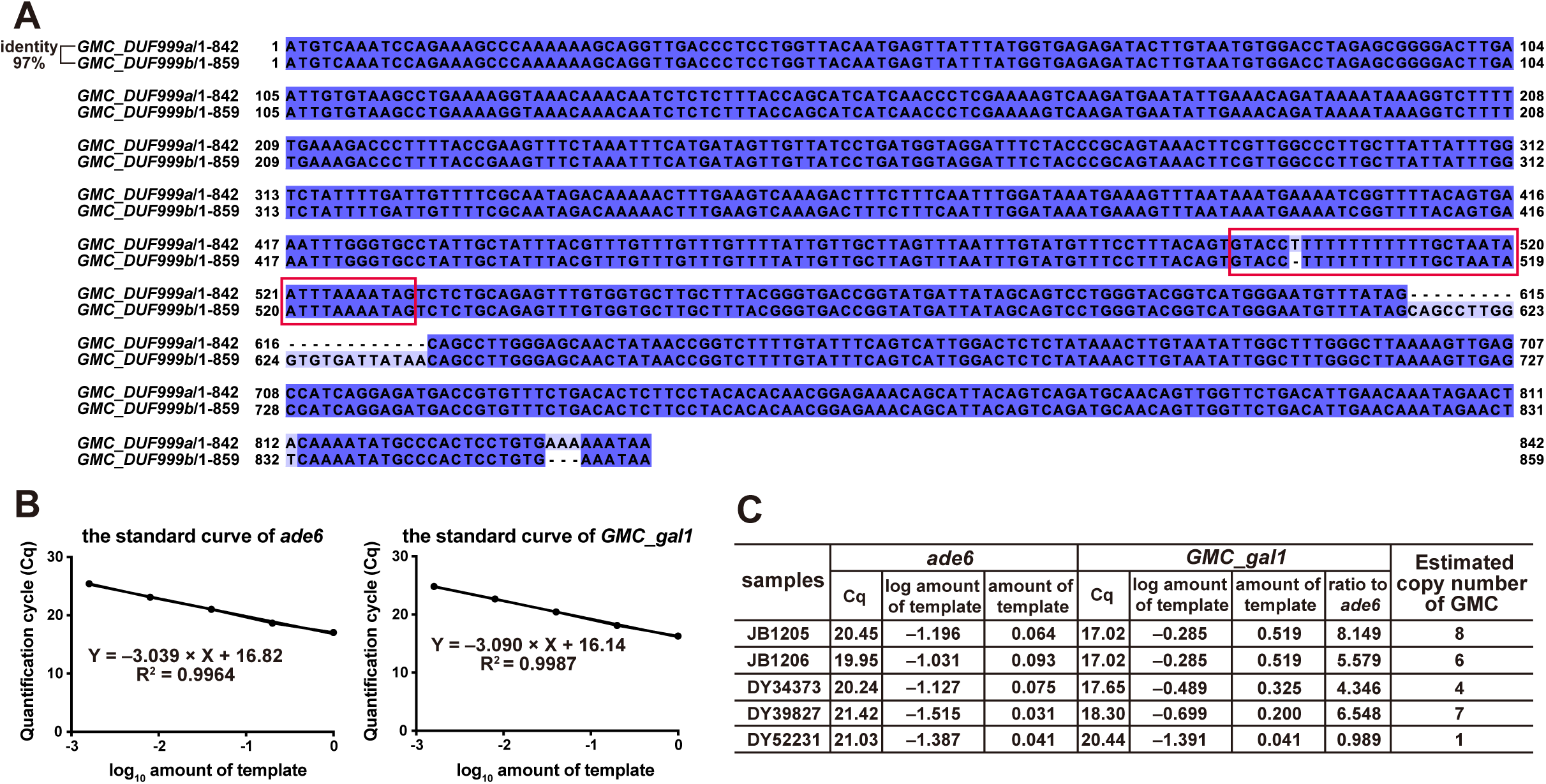
Sequence differences between *GMC_DUF999a* and *GMC_DUF999b* and copy number measurement of the GMC. (*A*) Sequence alignment of *GMC_DUF999a* and *GMC_DUF999b*, with identical bases highlighted using a dark blue background and the only intron indicated by a red box. (*B*) Standard curves for quantitative real-time PCR (qPCR) amplification of the reference gene *ade6* and the target gene *GMC_gal1*. (*C*) qPCR-based quantitative determination of GMC copy number. BLAST analysis against NGS-based assemblies revealed that the GMC is present in four of the 58 isolates: DY39827, DY34373, JB1205, and JB1206—the first two being the Gal^F^ isolates. DY52231 is a JB22-background strain containing a single copy of the *GMC_gal_region*.

**Fig. S9.**
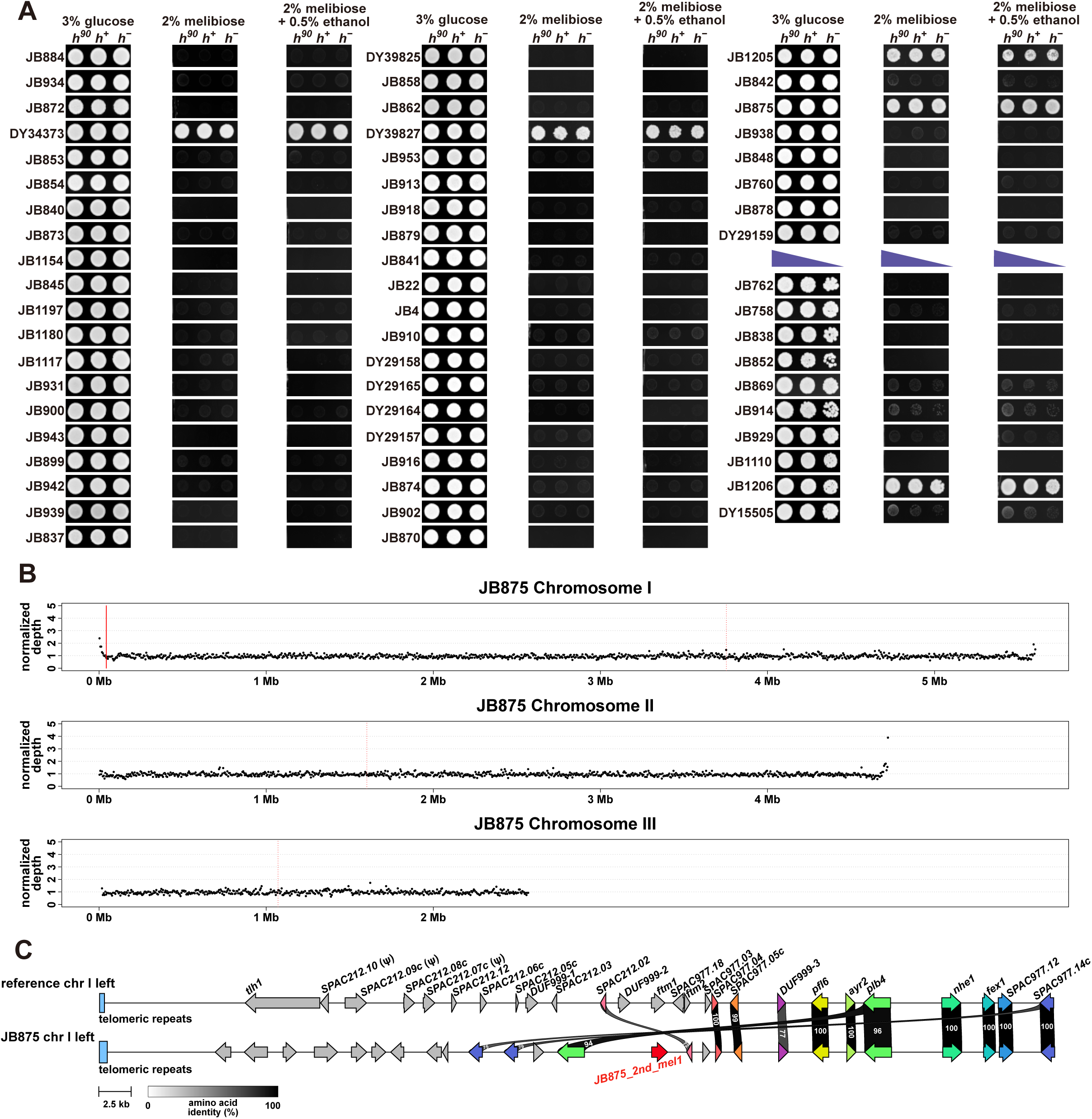
Analyses of melibiose utilization trait of 58 natural isolates and the copy number of the *JB875_2nd_mel1* gene in JB875. (*A*) Plate-based growth assay examining the melibiose utilization trait of 58 *S. pombe* natural isolates. Ethanol supplementation did not affect growth. For homothallic natural isolates (denoted as *h^90^* for brevity), the engineered heterothallic derivatives (*h^+^* and *h^−^*) were also examined. For natural isolates that cannot undergo sexual reproduction, multiple dilutions indicated by purple triangles were spotted on the plates. (*B*) Read depth plots showing the normalized depth of Illumina sequencing reads mapped to the PacBio HiFi-based JB875 assembly. Each black dot corresponds to a 5-kb window. The red solid line marks the position of *JB875_2nd_mel1*, while the red dotted lines indicate the locations of centromeres. (*C*) Synteny plot comparing the left-end regions of chromosome I in the reference genome and JB875. *JB875_2nd_mel1* is highlighted in red.

**Fig. S10.**
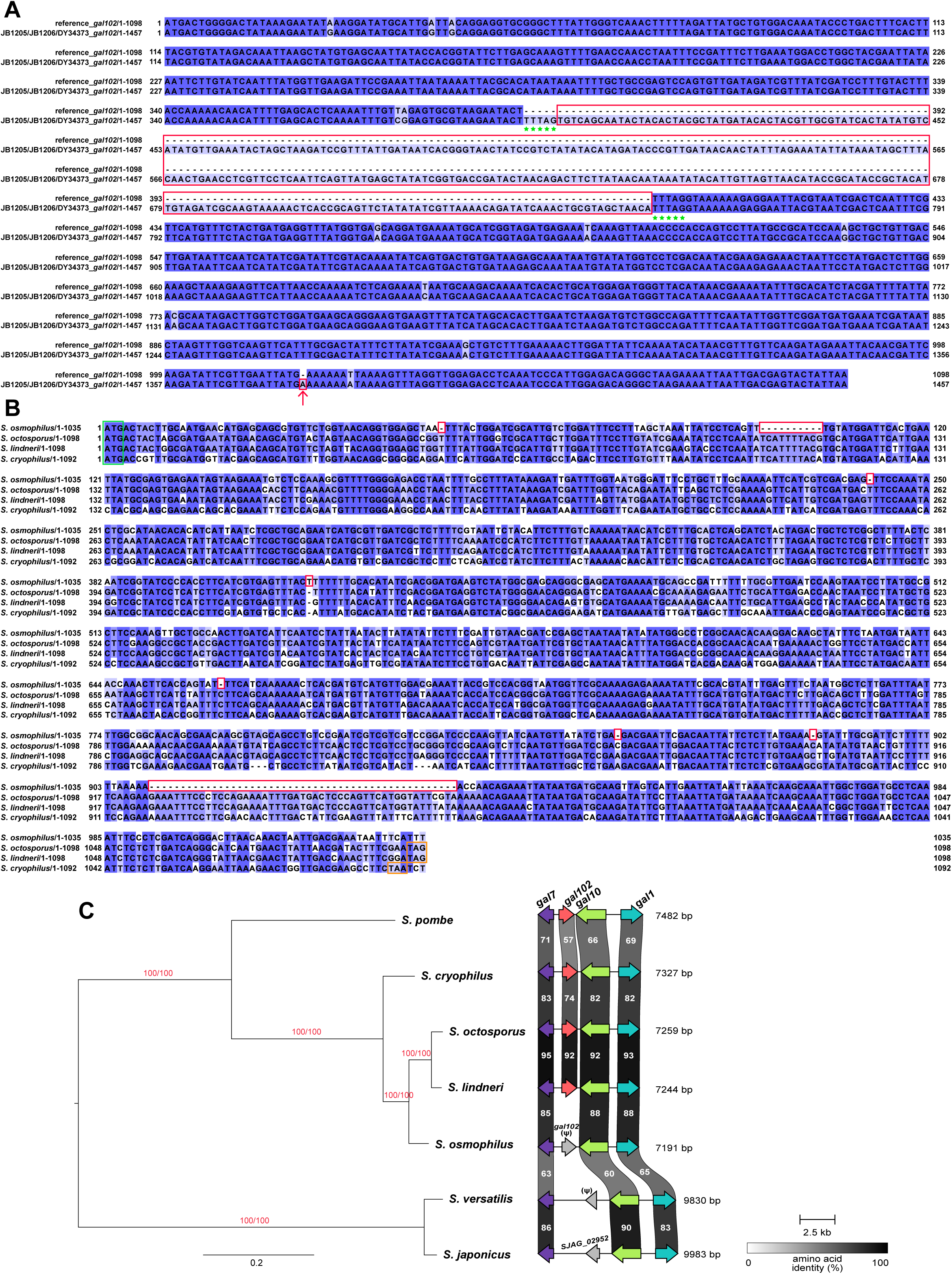
Multiple independent *gal102* inactivation or loss events. (*A*) Alignment comparing the coding sequences of the intact *gal102* gene in the *S. pombe* reference genome and the long terminal repeat (LTR)-containing *gal102* gene present in JB1205, JB1206, and DY34373. Identical bases are highlighted using blue background. The LTR is indicated by red rectangles. Green stars denote the 5-bp target site duplication (TSD) sequences (“TTTAG”) flanking the LTR. A frameshifting single-nucleotide insertion is denoted by a red box and an arrow. (*B*) Alignment comparing the coding sequences of the pseudogenized *gal102* gene in *S. osmophilus* and the intact *gal102* genes in *S. octosporus*, *S. lindneri*, and *S. cryophilus*. Identical bases are highlighted using blue background. Red boxes highlight frameshifting indels in the *S. osmophilus* sequence. Start codons are labeled with a green box and stop codons in *S. octosporus*, *S. lindneri*, and *S. cryophilus* sequences are marked with orange boxes. The *S. osmophilus* sequence lacks a stop codon at the expected position. (*C*) Maximum likelihood (ML) phylogeny and *gal* cluster synteny plot of fission yeast species. The ML phylogenetic tree on the left side was constructed using IQ-TREE and rooted by midpoint rooting. It was inferred with a concatenation matrix of 1128 complete and single-copy BUSCO genes present in 7 fission yeast species. Branch support values were calculated by IQ-TREE. The scale bar below tree indicates 0.2 substitutions per site. The symbol “Ψ” indicates a pseudogene. *S. japonicus* and *S. versatilis* lack *gal102*. In *S. japonicus*, the *SJAG_02952* gene situated between *gal7* and *gal10* encodes a nucleolar RNA-binding protein and is a paralog of *SJAG_00949*. In *S. versatilis*, a pseudogene sharing sequence similarity with *SJAG_02952* is located between *gal7* and *gal10*.

**Fig. S11.**
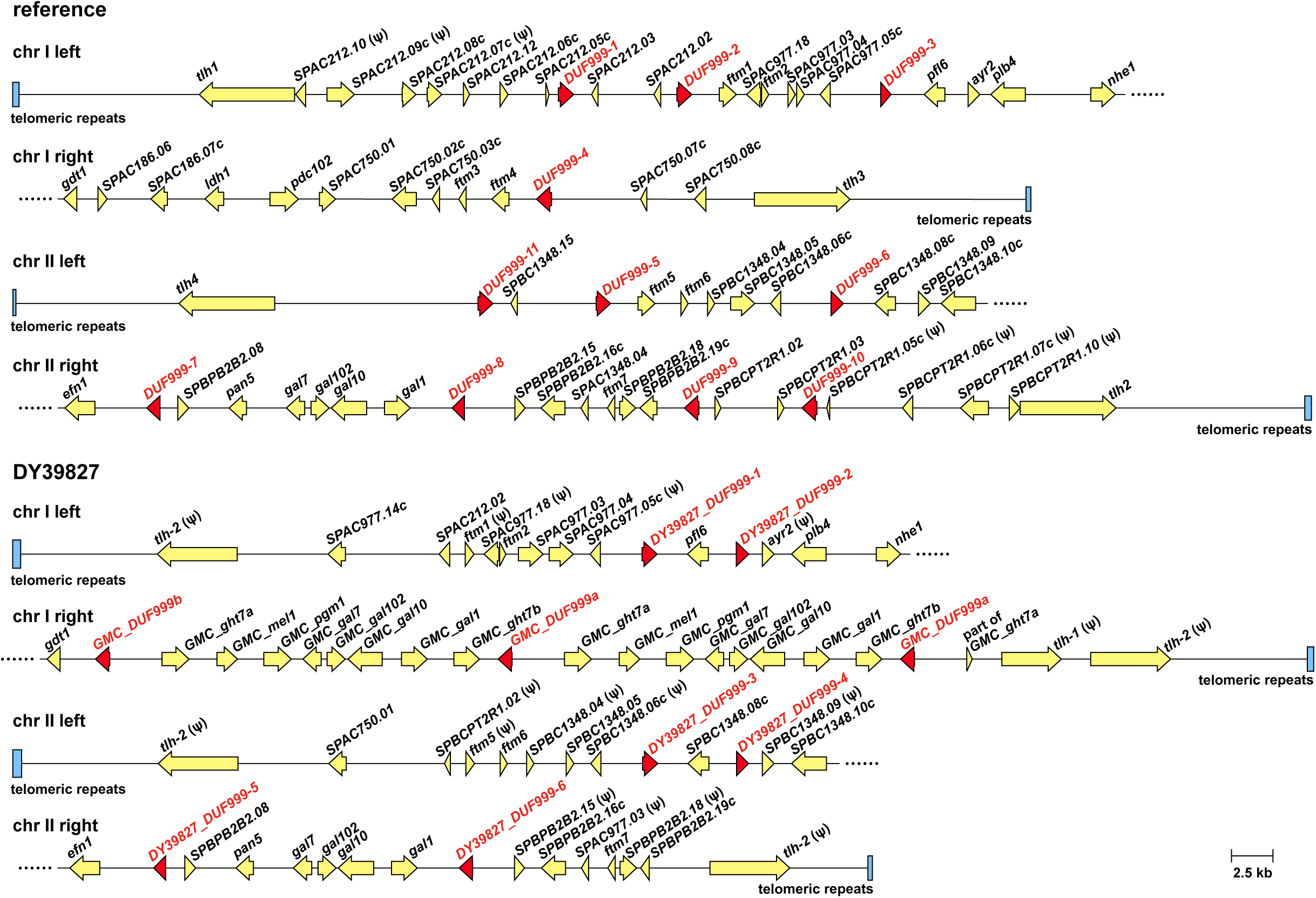
The genomic locations of *DUF999* genes in the *S. pombe* reference genome and the genome of DY39827. Schematic diagrams showing the left-end and right-end regions of chromosomes I and II in the *S. pombe* reference genome and the genome of DY39827. All *DUF999* genes are highlighted using red color. Except for *DUF999-11*, *DUF999* genes in the reference genome were numbered according to their chromosomal positions and the numbering is consistent with the numbering by PomBase. *DUF999-11* is a gene not annotated by PomBase. It is identical to *DUF999-10*. We numbered *DUF999* genes not associated with the GMCs in the DY39827 genome in the same way. Blue rectangles denote telomeric repeats.

**Fig. S12.**
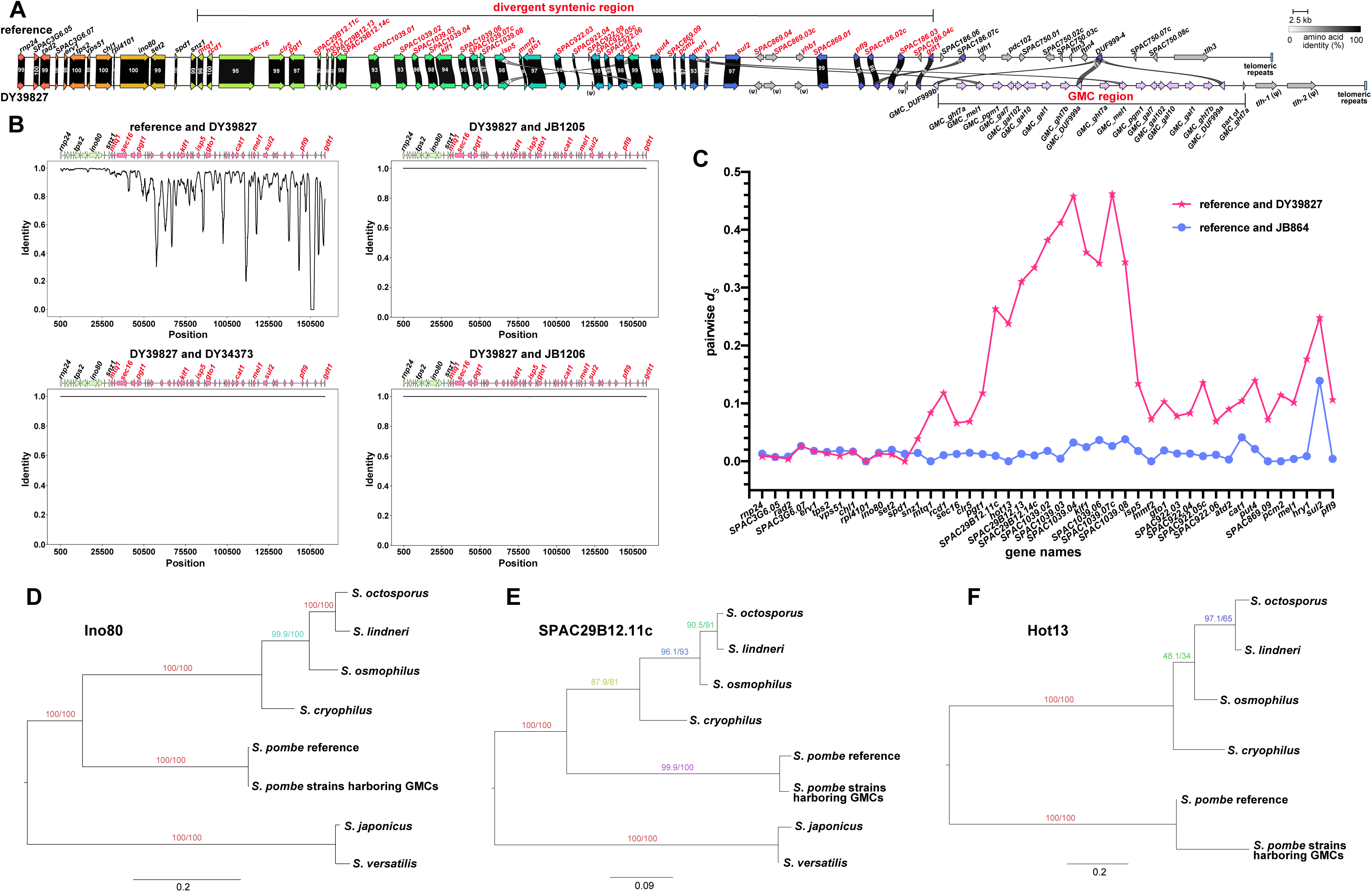
The divergent chromosome I subtelomeric region adjacent to the GMCs. (*A*) Synteny plot comparing the chromosome I right-end regions in the reference genome and the DY39827 genome. The divergent syntenic region and GMC region are highlighted using bracketed lines. The symbol “Ψ” indicates a pseudogene. (*B*) Sliding-window analysis showing the elevated nucleotide diversity between the reference genome and the DY39827 genome within the divergent syntenic region, which spans from the *mtq1* gene to the *gdt1* gene. Genes within the divergent syntenic region are colored pink in the diagram at the top. The four GMC-containing strains share nearly identical sequences in this region. The window size is 1 kb and the step size is 50 bp. (*C*) Line chart depicting pairwise *d_s_* values of genes located in the region shown in (*B*). The pink line denotes the *d_s_* values calculated based on the pairwise comparison between the reference strain and DY39827, and the blue line represents the *d_s_* values calculated based on the pairwise comparison between the reference strain and JB864, a representative of the NONREF strains. (*D-F*) Maximum likelihood (ML) phylogenetic trees of Ino80 and its homologs across fission yeasts (*D*), SPAC29B12.11c and its homologs across fission yeasts (*E*), Hot13 and its homologs across fission yeasts (*F*). The trees were constructed using IQ-TREE and rooted by midpoint rooting. Branch support values were calculated by IQ-TREE.

**Fig. S13.**
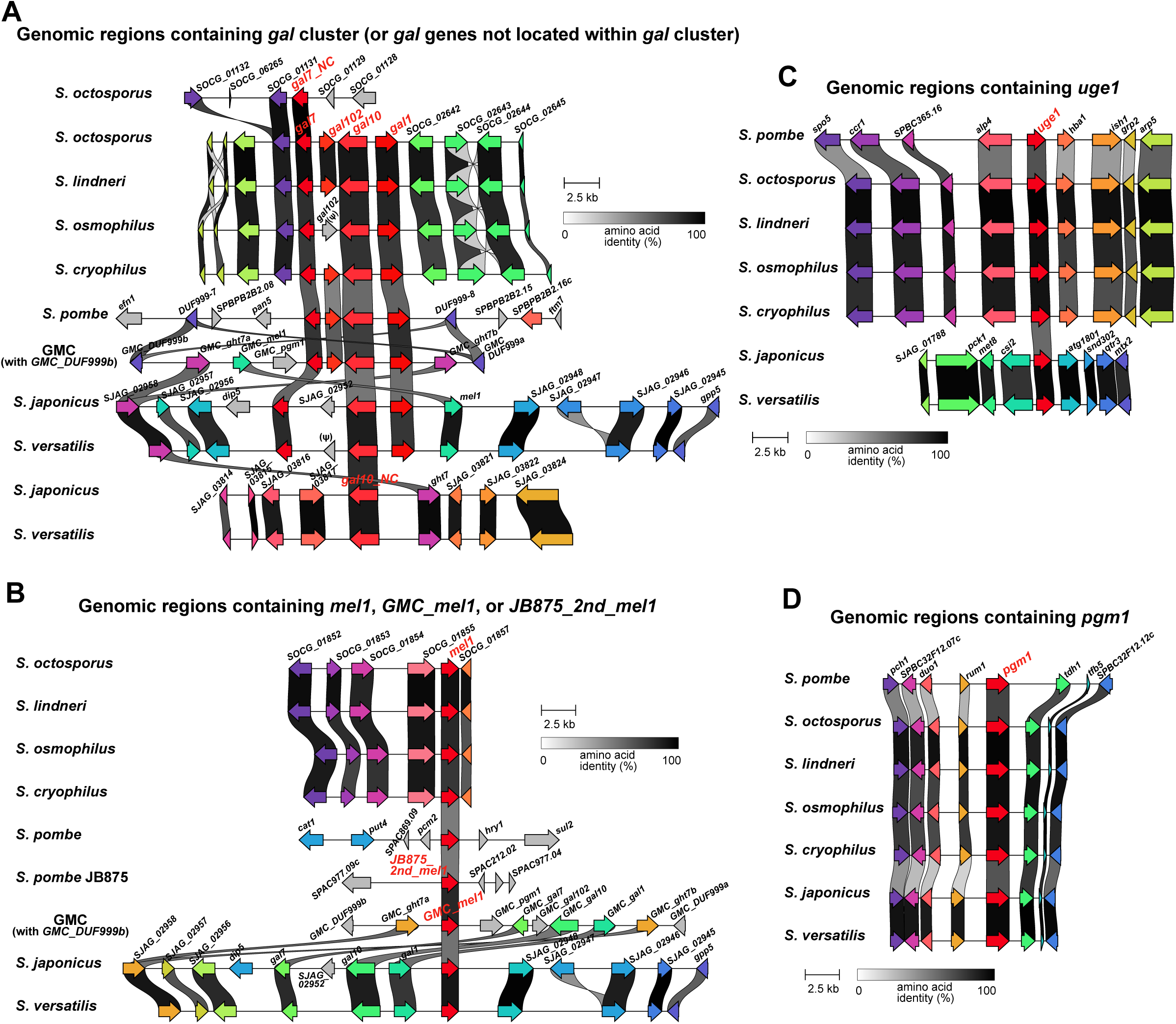
Cross-species synteny analysis of genomic regions containing galactose and melibiose utilization genes. (*A-D*) Synteny plots comparing genomic regions containing the *gal* cluster or *gal* genes not located within the *gal* cluster (*A*), *mel1*, *GMC_mel1*, or *JB875_2nd_mel1* (*B*), *uge1* (*C*), and *pgm1* (*D*). Galactose and melibiose utilization genes are highlighted in red. The appendix “NC” denotes Gal7- and Gal10-coding genes located outside the *gal* cluster.

